# Membrane lipid composition and endocytosis modulate Wingless release from secreting cells

**DOI:** 10.64898/2026.02.12.705585

**Authors:** Ines Alvarez-Rodrigo, David Willnow, Cyrille Alexandre, Claire Lebarbenchon, Annabel Borg, Heather Findlay, Ikram Arahouan, Yuguang Zhao, Florencia di Pietro, Svend Kjaer, Paula Booth, Reinhard Bauer, E. Yvonne Jones, Yohanns Bellaïche, Jean-Paul Vincent

## Abstract

Wnts are secreted signalling molecules that regulate development and adult homeostasis. Most Wnts carry a lipid moiety that must be shielded from the aqueous environment. In the secretory pathway, this is achieved by a hydrophobic tunnel in Wntless, a multipass transmembrane protein. However, the Wnt lipid moiety must be released from Wntless before Wnts can engage with Frizzled receptors on receiving cells. Here we address the cell biological basis of Wnt-Wntless dissociation, using as a model the secretion of Drosophila Wingless in wing primordia. Super-resolution microscopy shows that Wingless first reaches the apical surface before being re-internalized to reach, without Wntless, specialized Rab7/Rab4-positive endosomes. From there Wingless traffics to the basolateral membrane where it can engage with glypicans to form a basolateral gradient. Acute inhibition of endocytosis, either with a temperature-sensitive *dynamin* mutant or a novel optogenetic means of inhibiting clathrin, leads to apical Wingless release in abnormal punctae devoid of Wntless, suggesting that Wingless-Wntless dissociation commences at the apical surface, perhaps because of a distinct lipid composition there. Indeed, similar looking punctae are produced upon genetic abrogation of the ceramide synthase Schlank, specifically in Wingless-producing cells. These punctae resemble insoluble aggregates that form *in vitro* upon detergent removal. Accordingly, punctae formation can be prevented by shielding the Wingless lipid, *in vivo* with excess Dally-like protein (Dlp) or *in vitro* with liposomes. Our results show that membrane lipid composition modulates the orderly transfer of Wingless lipid from Wntless to the inner endosomal surface thus preventing aggregation and ensuring seamless secretion in the basolateral space.

## Introduction

Wnts are evolutionarily conserved signalling molecules with fundamental roles in stem cell maintenance, cell fate decisions and growth during development and adult homeostasis (Clevers et al., 2014; Logan and Nusse, 2004; Loh et al., 2016; Nusse and Clevers, 2017; Steinhart and Angers, 2018). All characterised Wnt proteins (except arthropod WntD) carry a palmitoleoyl moiety, a monounsaturated 16-carbon fatty acid appended to a conserved serine residue in the ER by the O-acyl transferase Porcupine (Ching et al., 2008; Herr and Basler, 2012; Murat et al., 2010; Takada et al., 2006). The lipid moiety of Wnts mediates their interaction to Frizzled receptors (Hirai et al., 2019; Janda et al., 2012; Nile et al., 2017; Tsutsumi et al., 2020). As expected therefore, “unlipidatable” Wnt proteins created by mutating the conserved acylated serine have strongly reduced signalling activity (Franch-Marro et al., 2008; Luz et al., 2014; Speer et al., 2019; Tang et al., 2012). While the lipid moiety is essential for signalling activity of Wnts, it substantially reduces their solubility (Takada et al., 2006; Willert et al., 2003), raising a challenge for secretion and release by producing cells.

In the secretory pathway, the solubility challenge is met by Wntless (Wls), a multi-pass transmembrane protein that escorts lipidated Wnts from the ER to the cell surface. In the absence of Wls, Wnt proteins remain trapped within expressing cells (Bänziger et al., 2006; Bartscherer et al., 2006; Goodman et al., 2006; Gross et al., 2012; Moti et al., 2019; Najdi et al., 2012; Yu et al., 2014). Wls shields the Wnt lipid in the secretory pathway by virtue of a hydrophobic tunnel formed by transmembrane helices (Nygaard et al., 2021; Qi et al., 2023; Zhong et al., 2021). Since Wls is a multipass transmembrane protein, it has been suggested that Wnts could be released as a complex with Wls on exosomes or non-exosomal high density particles (Beckett et al., 2013; Chen et al., 2016; Gross et al., 2012; Koles et al., 2012; Korkut et al., 2009; Yang et al., 2025). Wnt-Wls complexes have also been seen on cytonemes, which could also conceivably account for the spread of Wnts (and Wls) beyond expressing cells (Piers et al., 2024). However, evidence from Drosophila wing imaginal discs shows that Wingless, the main Drosophila Wnt, is released from secreting cells without Wls, implying dissociation before release (Beckett et al., 2013; Gross et al., 2012; McGough et al., 2020). Indeed, a number of proteins, including secreted Frizzled-related proteins (sFRPs), Afamin, WIF1 and Dlp class glypicans have been suggested to accommodate the Wnt lipid after dissociation from Wls (de Almeida Magalhaes et al., 2024; Mihara et al., 2016). Whether Wnts are released with or without Wls from producing cells, their lipid must ultimately disengage from Wls’ hydrophobic tunnel so that it can bind to a Frizzled receptor to trigger signalling. Therefore, a mechanism must exist to trigger the release of Wnts from Wls after their joint progression through the secretory pathway.

Recent studies have elucidated the structural basis of Wnt-Wls complex formation (Nygaard et al., 2021; Qi et al., 2023; Zhong et al., 2021). However, the mechanism that regulates dissociation for Wnt release is still largely unknown. Vacuolar pH decreases progressively from the ER to secretory vesicles, suggesting that vacuolar acidification could trigger dissociation. Accordingly, vATPase inhibitors prevent Wnt3a secretion from cultured cells (Coombs et al., 2010). However, lowering the pH is not sufficient to dissociate the Wnt-Wls complex in immunoprecipitates (Coombs et al., 2010), raising the possibility that another mechanism could be at work. Another characteristic of the secretory pathway is that its lipid composition varies with maturation. Thus, high concentration of phosphatidylcholine found in the ER could promote Wnt binding to Wls in the ER (Qi et al., 2023), while a different lipid composition at the plasma membrane could trigger release. Alternatively, dissociation could be triggered by the change in membrane curvature expected to occur when secretory vesicles fuse with the plasma membrane (Alvarez-Rodrigo et al., 2023). It has been suggested that, in addition or in parallel, the glycosylation pattern, which can differ among Wnts, could affect the strength of the Wnt-Wls complex beyond the Golgi (Yamamoto et al., 2013). Finally, a protein resident of the secretory pathway, e.g. TMEM132A, could help stabilize the Wnt-Wls complex specifically in the secretory pathway (Li and Niswander, 2020). Overall, several mechanisms have been proposed to trigger Wnt release from Wls but so far, their physiological relevance in vivo remains to be assessed.

In the developing wing of Drosophila, Wg is produced as a thin stripe at the dorsal-ventral boundary and forms a basolateral gradient to modulate growth. In earlier work we have shown that Wg is first targeted to the apical surface of Wg-producing cells before being endocytosed back into the same cells for basolateral release. We have also shown that such Wg transcytosis requires the ubiquitin ligase Godzilla and its target the SNARE VAMP3/Synaptobrevin (Yamazaki et al., 2016). More recently, the class III phosphatidylinositol 3-kinase (PI3K(III)) complex was also shown to be involved (Holzem et al., 2025). Disrupting Wg transcytosis results in abnormal (sub)apical secretion (Gross et al., 2012; Linnemannstons et al., 2020; Sharma et al., 2026; Witte et al., 2020) suggesting that transcytosis is important for orderly Wg production within the developing wing.

In the present study, we first combine super-resolution microscopy with the genetic toolbox of Drosophila to track the fate of Wg and Wls after they have trafficked together to the apical plasma membrane. We show that, upon acute inhibition of endocytosis, Wg is released abnormally as apical punctae, and unable to form a basolateral gradient. From this observation, we infer that Wg-Wls dissociation commences at the apical plasma membrane, but that rapid endocytosis ensures that most Wingless is re-internalised and retained by producing cells. Dissociation could continue within endocytic pits but is completed soon thereafter since Wg is no longer colocalising with Wls when it reaches characteristic endocytic vesicles marked by Rab4 and/or Rab7. From there, Wingless can continue its journey to the basolateral membrane for release. Super-resolution microscopy shows that Wg lines the inner surface of endocytic vesicles, suggesting membrane association. We provide evidence that this association does not require Dlp and that the Wg lipid can insert directly into the lipid bilayer of liposomes in vitro. Knocking-down the ceramide synthase Schlank triggers premature release from the membrane and the formation of aggregate-like punctae, suggesting that maintaining the right balance of ceramide-derived lipids (or their precursors) is needed for orderly insertion of the Wnt lipid in the bilayer and transcytosis to the basolateral surface where the Wingless lipid could insert in the hydrophobic tunnel of Dlp for further transport.

## Results

### Wg traffics to Rab4/7-positive vesicles after dissociation from Wls

Fixed wing imaginal discs stained with anti-Wg antibodies were imaged with super-resolution microscopy by optical re-assignment (SoRa) followed by deconvolution (see Materials and Methods). This revealed the existence, within Wg-producing cells, of vesicle-like structures enriched with Wg protein at their periphery (Fig1A). Similar structures could be recognised with GFP fluorescence in unfixed embryos expressing tagged GFP-Wg from the endogenous locus (Port et al., 2014) (Fig1B, FigS1A), excluding the possibility of a fixation artifact. These structures could not be resolved with conventional confocal microscopy but were visible with other super-resolution microscopy techniques (FigS1B), indicating that they are not a deconvolution artifact. They were also recognisable in imaginal discs endogenously expressing SNAP-Wg labelled with a permeable SNAP-tag, but not with an impermeable SNAP-tag without permeabilization, suggesting that they are vesicular in nature and that Wg molecules are associated with their inner membrane (FigS1C). We therefore term these structures Wg-lined vesicles (WgLVs) and devised an image analysis pipeline based on radial fluorescence for their rapid detection (FigS1D-H). The abundance of Wg at WgLVs contrasts with the lack of Wls enrichment (Fig1C). This is not due to differential accessibility of antibodies to intracellular compartments since the two antibodies showed Wg and Wls colocalising both in the cis-Golgi (identified by the presence of GMAP; (Friggi-Grelin et al., 2006)) (Fig1D-E), and at the apical-most region of Wg-producing cells (Fig1F). Of note, a Wls-GFP fusion protein (*wls[ExGFP]*; (Yu et al., 2020)) that partially retains Wg in secreting cells, perhaps because the GFP moiety interferes with dissociation, becomes detectable at WgLVs (FigS2). Conversely, an endogenously expressed lipidation mutant Wg (Wg^S239A^; (Franch-Marro et al., 2008; McGough et al., 2020; Takada et al., 2006)), which traffics independently of Wls to the plasma membrane, does not accumulate in characteristic vesicles (Fig1G-H). We conclude that Wg normally travels with Wls through the secretory pathway, but these proteins normally dissociate before Wg reaches WgLVs.

**Figure 1.**
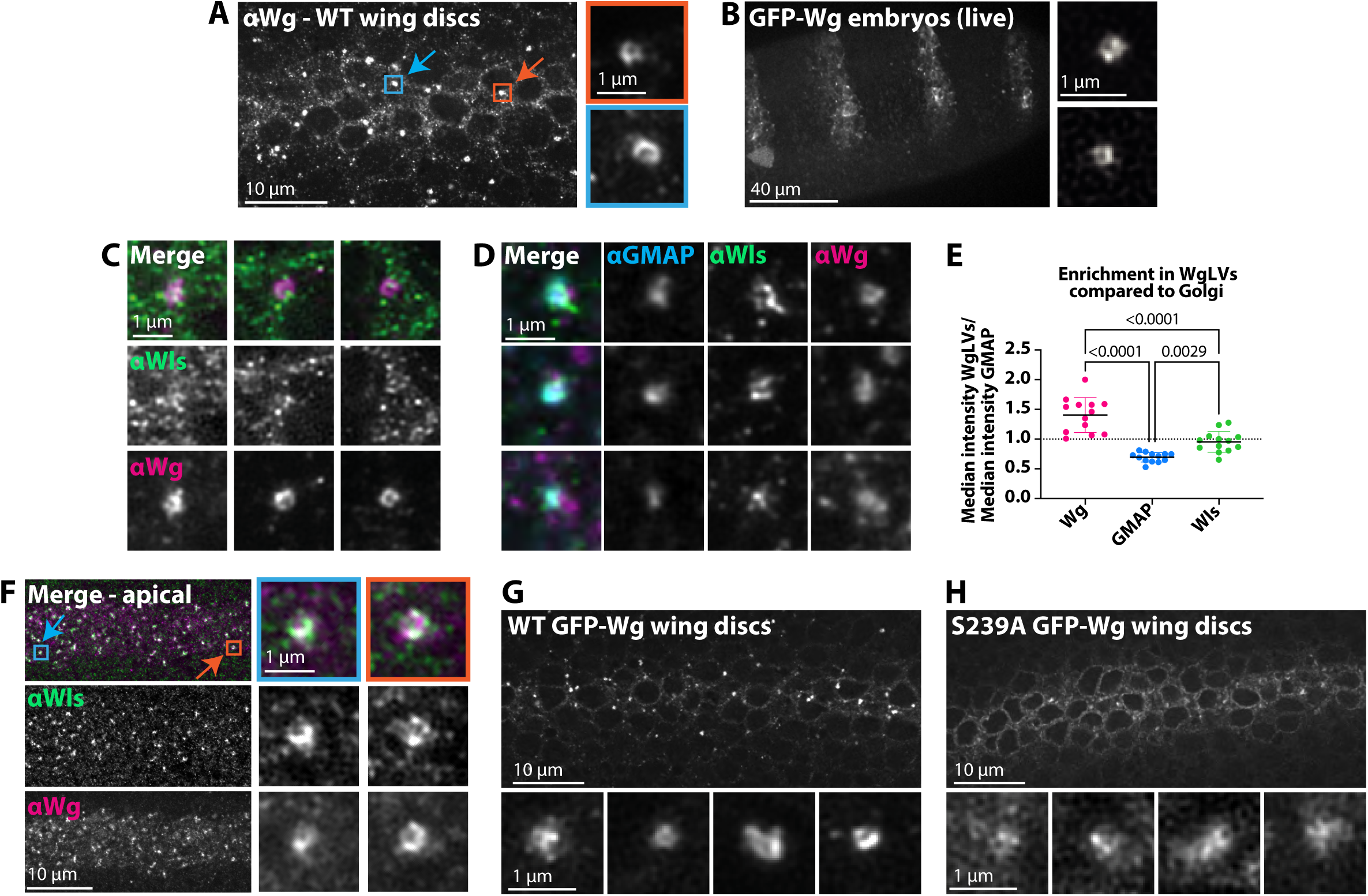
Lipidated Wg is enriched in Wg-lined vesicles (WgLVs) without Wls. **(A)** Total Wg staining of Wg-producing cells in wild-type wing discs imaged with super-resolution microscopy. In the middle of the apical-basal axis, Wg is particularly concentrated in bright vesicles, two of which are highlighted with *blue* and *orange* boxes. Insets on the right (with adjusted display settings), show Wg enrichment at the periphery of the vesicles. **(B)** Representative super-resolution image from a live Drosophila embryo expressing GFP-Wg from the *wg* locus. **(C)** Examples of WgLVs in wild-type wing discs co-stained for Wls. While Wg is enriched in the periphery of WgLVs, Wls is not. **(D)** By contrast, Wg and Wls are both found in GMAP-positive vesicles. **(E)** Quantification of the enrichment of Wg, GMAP or Wls in WgLVs, as compared to that in Golgi. N = 13 wing discs from three independent experiments, with >80 WgLVs and >100 GMAP+ particles analysed per disc. Statistical test used: paired one-way ANOVA (comparing WgLV to Golgi signals from the same image) with Geisser-Greenhouse correction and Tukey’s multiple comparisons test. Graph bars: mean with SD. **(F)** Apical-most region of Wg-producing cells, with examples of vesicles (possibly endocytic pits) where Wg and Wls co-localize. **(G-H)** Enrichment of wild-type GFP-Wg but not unlipidatable GFP-Wg^S239A^ (both expressed from the *wg* locus) in WgLVs.

To identify the nature of WgLVs, we stained a panel of wing discs expressing endogenously tagged endocytic markers with anti-Wg. Rab4, a marker of fast recycling endosome (De Renzis et al., 2002) and Rab7, which marks late endosomes (Rink et al., 2005) were both found to colocalise with Wg at WgLVs (Fig2A-B and FigS3A). The average radial fluorescence profile from YFP and anti-Wg in YFP-Rab4- or YFP-Rab7-expressing discs showed that both markers appear to surround WgLVs (Fig2C). We infer therefore that Wg lines the inner membrane of WgLVs while Rab4 and/or Rab7 coat the outside (FigS3B). One should note that not all WgLVs were labelled by YFP-Rab4 or YFP-Rab7: 50% of WgLVs were YFP-Rab4-positive and 75% were YFP-Rab7-positive (FigS3C-F). They tended to be larger and to contain more Wg than Rab4/7-negative WgLVs (FigS3C-D). They were also most abundant in the mid-lateral region of Wg secreting cells (FigS3E-F). Many WgLVs were labelled by both YFP-Rab4 and anti-Rab7 antibodies (Fig2D-E), two markers that are not known to normally colocalise. These double-positive vesicles must nevertheless be genuine mature endosomes and not lysosomes since they colocalised with TagRFP-Vps35, an endocytic marker (Korolchuk et al., 2007; Seaman et al., 2009). Although a time course of the progression of WgLVs in the endocytic pathway cannot be directly observed, their distribution along the apical-basal axis is consistent with a progression from Rab4/7 double negative (apically), to double positive (laterally) and finally a mixture of different identities in the basal region (FigS3E-F). Overall, super-resolution imaging suggests that Wg and Wls have already dissociated when they reach endocytic vesicles associated to various degrees with Rab4 and Rab7.

**Figure 2.**
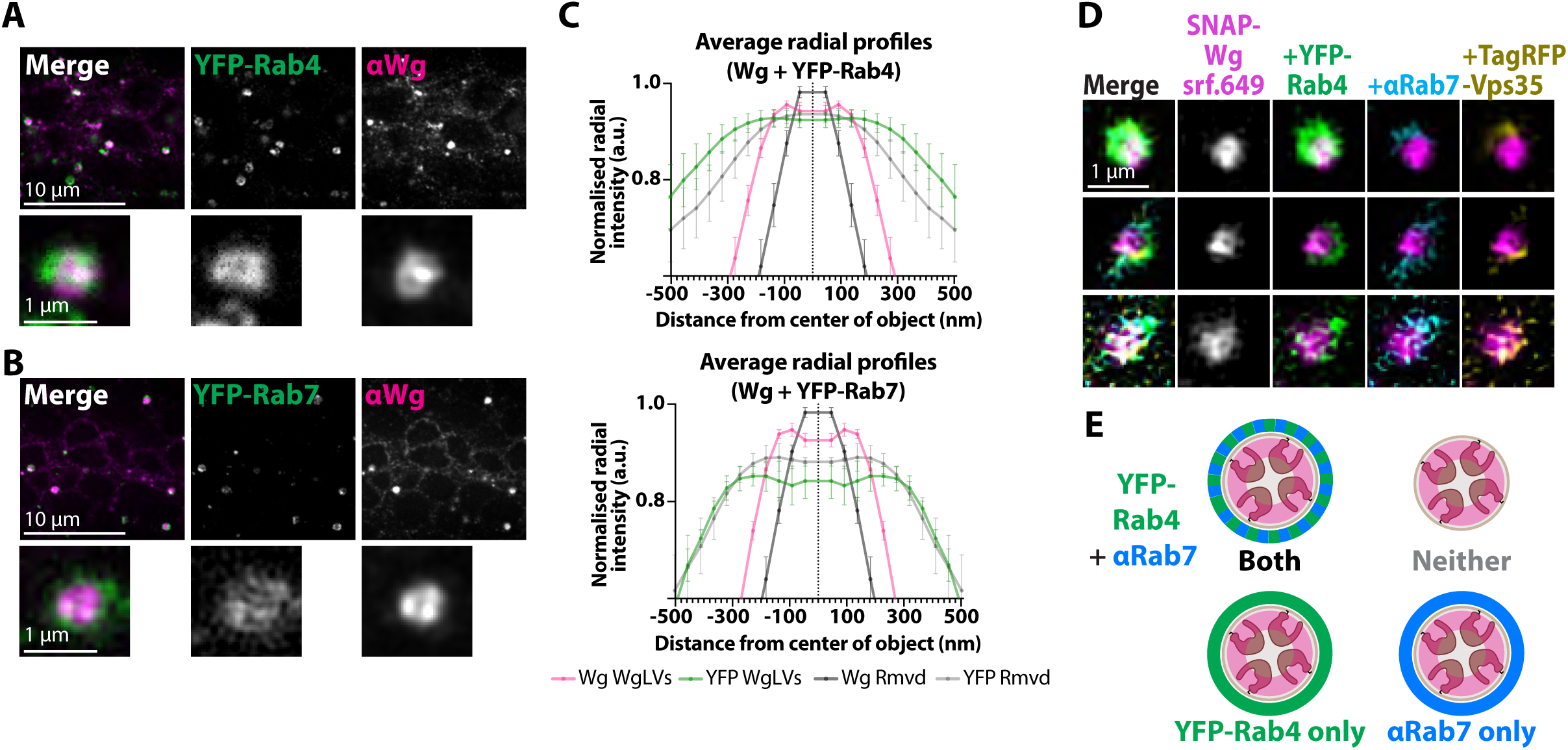
WgLVs are Rab4 and Rab7 positive vesicles. (A-B) Wing discs endogenously expressing either YFP-Rab4 *(A)* or YFP-Rab7 *(B)* and stained for Wg. One example of a colocalising WgLV is shown for each Rab. (C) Average radial fluorescence profiles of YFP-Rab4-positive (N= 13 wing discs) or YFP-Rab7-positive (N = 4 wing discs) WgLVs. Error bars indicate SD. *Magenta* indicates the Wg profile, and *green* the corresponding Rab. In *grey* and *light grey* we show the profiles of bright Wg-containing particles that did not pass our pipeline to identify WgLVs and hence removed from the WgLV plots (“Rmvd”). (D) Examples of WgLVs (*magenta/grayscale*) positive for YFP-Rab4 (*green*), anti-Rab7 (*blue*) and TagRFP-Vps35 (*yellow*). (E) Diagram illustrating the different populations of WgLVs in terms of Rab4/7 colocalization.

### Inhibition of endocytosis leads to abnormal apical release of Wg

Wls escorts Wingless through the secretory pathway all the way to the apical plasma membrane (Sharma and Chaudhary, 2024; Yu et al., 2014). Therefore, these two proteins must dissociate after this point, but before Wg is trafficked to Rab4/7-positive WgLVs. To assess whether release can occur upon arrival at the apical basal membrane, we looked in detail at the distribution of Wg artificially held at the plasma membrane by acute inhibition of endocytosis with a temperature-sensitive Dynamin encoded by *shibire^ts^.* In this genotype, endocytosis proceeds normally at permissive temperature (18°C), but endocytic pits cannot progress into early endosomes at the restricted temperature (34°C). This phenotype is rapidly induced and reversible, making *shibire^ts^* a particularly powerful tool to manipulate endocytosis (Koenig and Ikeda, 1989). Previous work from our group showed that culturing *shibire^ts^* larvae at 34°C for 30-60 min is sufficient to cause Wg to accumulate at the extracellular side of the apical plasma membrane (Yamazaki et al., 2016) (Fig3A), while basolateral trafficking resumes rapidly upon return to permissive temperature. To track the progression of Wingless to WgLVs after resumption of endocytosis, we turned to super-resolution microscopy.

Total Wg protein was imaged in five conditions: control wild-type discs (*1*), control *shibire^ts^* at 18°C (*5*), *shibire^ts^* at 34°C for one hour (*2*), and *shibire^ts^* cultured for 15 (*3*) or 30 min (*4*) at 18°C after a 1-hour block at 34°C (Fig3B). In *shibire^ts^* imaginal discs raised at 18°C, Wg was homogeneously distributed at the apical surface of Wg-producing cells and was found in bright WgLVs in the lateral region, as is the case in wild type discs (Fig3C, *1* and *5*). Upon inhibition of Dynamin at 34°C, Wg accumulated at the apical plasma membrane of secreting cells, with the appearance of numerous abnormal bright Wg punctae (Fig3C, *2*). Accumulation of Wg at the apical surface was accompanied by a marked reduction in the number of WgLVs in the lateral and basal regions, confirming that the supply route to WgLVs has been cut off. After return to permissive temperature, WgLVs reappeared, initially larger in size and brighter than steady state WgLVs (Fig3D). Within 30 min, the apical punctae had disappeared (Fig3C, *3* and *4*) indicating a return to homeostasis. Of note, no Wls was detected with the apical Wg punctae in neighbouring cells (Fig3E), suggesting Wg release without Wls (Fig3F).

**Figure 3.**
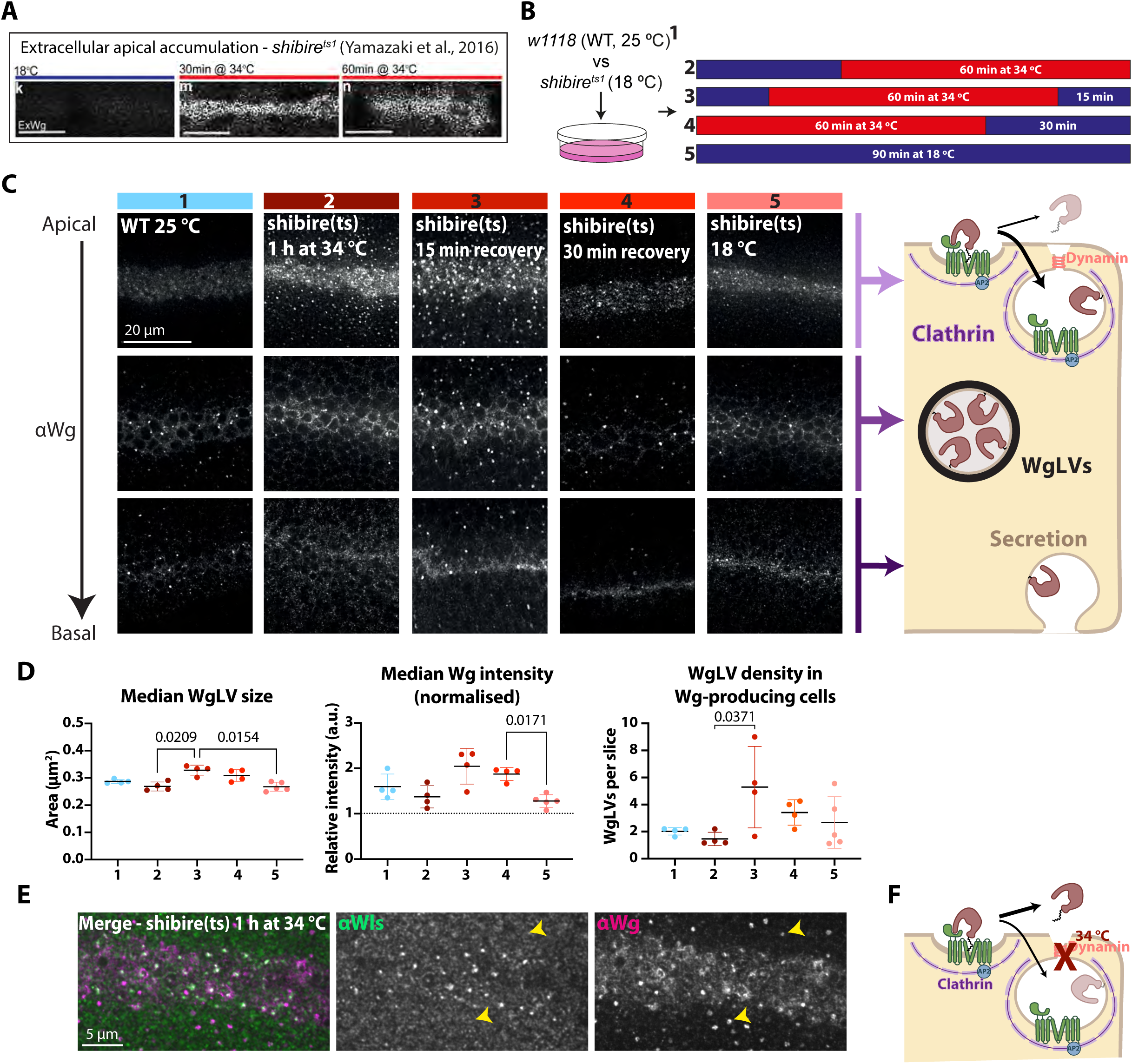
Wg dissociates from Wls during endocytosis before accumulating in WgLVs. **(A)** Effect of blocking endocytosis with *shibire^ts^* on extracellular Wg, adapted from (Yamazaki et al., 2016). Scale bar = 24 μm. **(B)** Experimental protocol to track Wg trafficking from the plasma membrane to WgLVs. **(C)** Representative super-resolution images (optical sections) of wing discs from each experimental condition (columns *1 – 5*). The diagram illustrates Wg progression from the apical to basal surface during transcytosis. **(D)** WgLV parameters extracted from the data exemplified in *(C)*. For each disc, we calculated a median WgLV size, median Wg intensity (normalised to total Wg intensity at the DV boundary) and WgLV density (number of WgLVs within Wg-producing cells in the whole *z-*stack, divided by number of images in the stack). >100 WgLVs analysed per disc. Graph shows mean and SD of the calculated values for N=4 discs (conditions *1 – 4) or* N= 5 discs (condition *5)*. Statistical test: one-way ANOVA with Welch correction and Dunnett’s T3 multiple comparisons test. **(E)** Apical-most region of a *shibire^ts^* wing disc kept a restrictive temperature for 60 min. Within Wg-producing cells, Wg and Wls colocalise, while in the receiving neighbours, Wg accumulate in abnormal bright punctae without Wls (*yellow arrowheads*). **(F)** Interpretation of results: inhibition of dynamin prevents closure of endocytic pits allowing content to leak out. While some Wg molecules are retained by Wg-producing cells, others are released without Wls and allowed to reach adjacent cells.

To further test if inhibition of endocytosis elicits apical Wg release, we developed optogenetic means of inactivating Clathrin. First, we generated a strain expressing functional GFP-tagged Clathrin heavy chain (GFP-Chc) from the endogenous locus (Fig4A-B) (see Materials). The GFP moiety was then used to trigger knock sideway or oligomerisation. For knock sideway, we used a light-inducible GFP trap based on the Magnet system (Kawano et al., 2015), with nMag-antiGFP nanobody and CD8-pMag each expressed from a Gal4-dependent transgene (Fig4C). Upon blue light illumination, the nMag and pMag moieties dimerise and drag GFP-Chc away from its function at coated pits. In discs expressing the trap components with *wg-Gal4*, but kept in the dark, the distribution of Wg, including at the apical surface, appeared normal. However, exposure to blue light for 1.5 h led to the formation of apical Wg punctae (Fig4C) similar to those seen after Dynamin inactivation (Fig3C). To complement the knock sideway approach, we adapted the Cry2olig approach which had been used to trigger light-inducible oligomerisation to acutely inhibit Clathrin in cells (Taslimi et al., 2014). However, instead of directly tethering Cr2olig and Chc, a *ubi*-Cry2oligVHH-mCherry transgene was introduced in the GFP-Chc background to oligomerise GFP-Chc. Light-mediated Cry2olig oligomerisation, as verified by the distribution of mCherry fluorescence, led to clustering of GFP-Chc, and this was associated with the accumulation of abnormal Wg punctae in the apical region (Fig4D). We conclude that inhibition of endocytosis elicits Wg release from the apical membrane. We suggest that, under normal circumstances, release from the apical plasma membrane is a slow process that is only readily detected if Wg is allowed to linger there.

**Figure 4.**
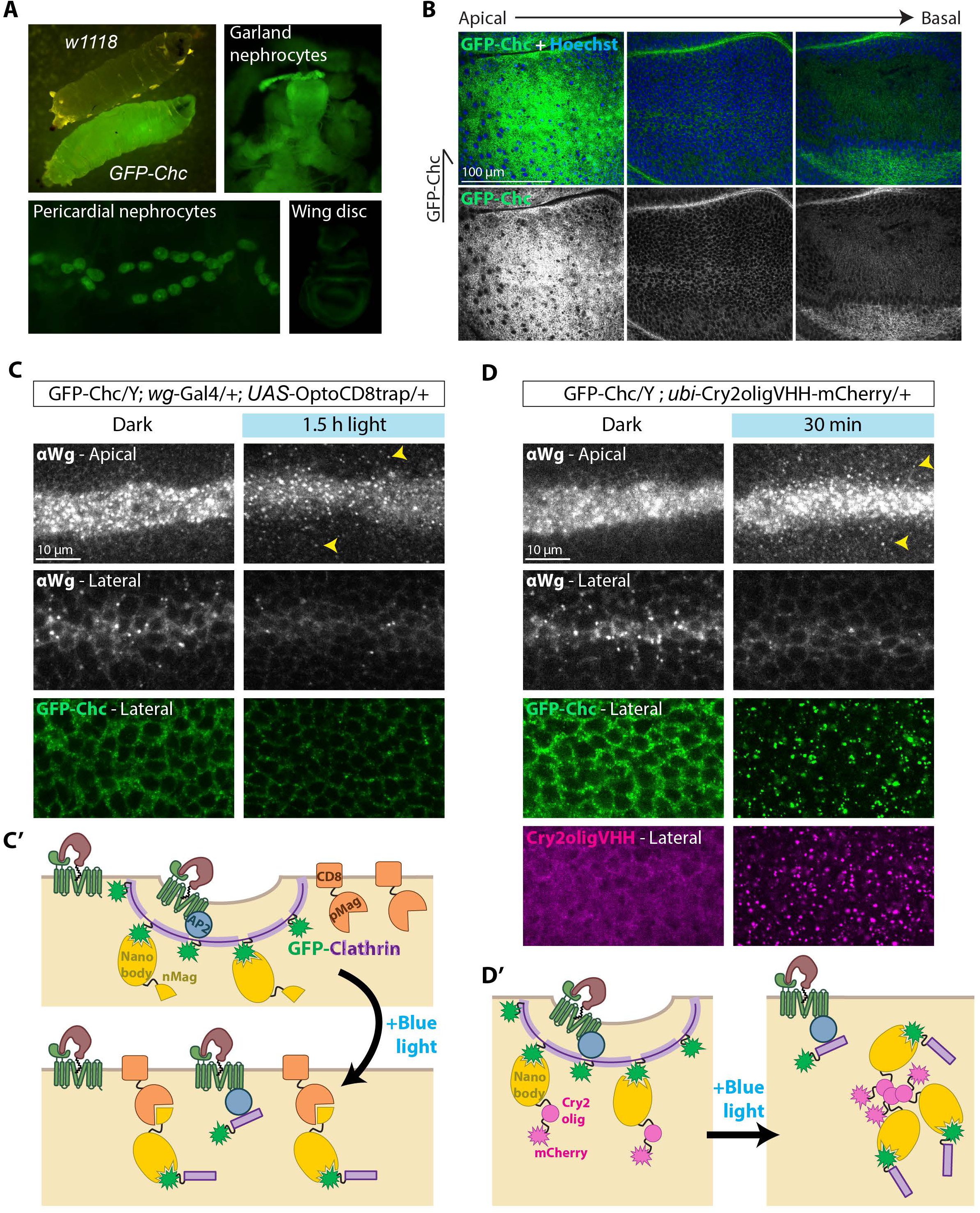
Light-induced trapping of endogenous GFP-tagged clathrin triggers Wg release from the apical membrane. (A) *GFP-chc* larvae are homozygous viable and widely express GFP. Garland and pericardial nephrocytes are particularly bright compared to other tissues such as wing discs. (B) Optical sections of fixed hemizygous *GFP-chc* wing discs. (C) Distribution of total Wg and GFP-Chc after optogenetic trapping of Chc. Hemizygous *GFP-Chc* males (all Chc is tagged) were used in combination with *wg-Gal4* and *UAS-OptoCD8trap*. Larvae maintained in the dark were used as control. Chc inactivation leads to apical release of abnormal Wg punctae (*yellow arrowheads*) and depletion of WgLVs. Total basal Wg seems unaffected (not shown). As the trap is at the membrane, GFP-Chc distribution appears normal, but fainter. (D) Distribution of total Wg and GFP-Chc after optogenetic clustering of GFP-Chc with ubiquitously expressed Cry2olig-VHH-cherry. Clustering is readily seen in the cherry and GFP channels. As with the optotrap, Chc inactivation leads to formation of apical Wg punctae and depletion of lateral WgLVs.

### Ceramide synthase knockdown triggers premature Wg release and accumulation in abnormal punctae

Because of their lipid adduct, Wnts are intimately associated with membranes, either indirectly through interaction with membrane-associated proteins such as Wls, or perhaps directly, as suggested by loading on artificial liposomes (Janeczek et al., 2017; Morrell et al., 2008; Tüysüz et al., 2017). Considering the relevance of lipids to Wnt trafficking, we revisited an early report that knockdown of *schlank*, which encodes the only Drosophila ceramide synthase, causes the accumulation of Wg punctae and reduces Wg signalling (Pepperl et al., 2013). We confirmed the formation of abnormal punctae by conventional confocal microscopy of imaginal discs expressing a *schlank* RNAi transgene (Fig5A). We also confirmed with *naked-NeonGreen*, a reporter of Wg signalling, that *schlank* RNAi reduces Wg signalling, despite the accumulation of Wg. Super-resolution microscopy revealed the presence of both WgLVs and large punctae lacking a visible lumen in *schlank*-deficient Wg-producing cells. Neither structure appeared to retain Wls (Fig5A’-A’’). Of note, abnormal punctae could be seen throughout the apical-basal axis, unlike the punctae of endocytosis-deficient cells (see below). A similar phenotype was seen upon inactivation of a conditional *schlank* knockout allele (*FRT-schlank-HA-FRT*) (FigS4A-B). We conclude that reduction of ceramide synthesis interferes with Wg targeting to WgLVs as well as signalling.

**Figure 5.**
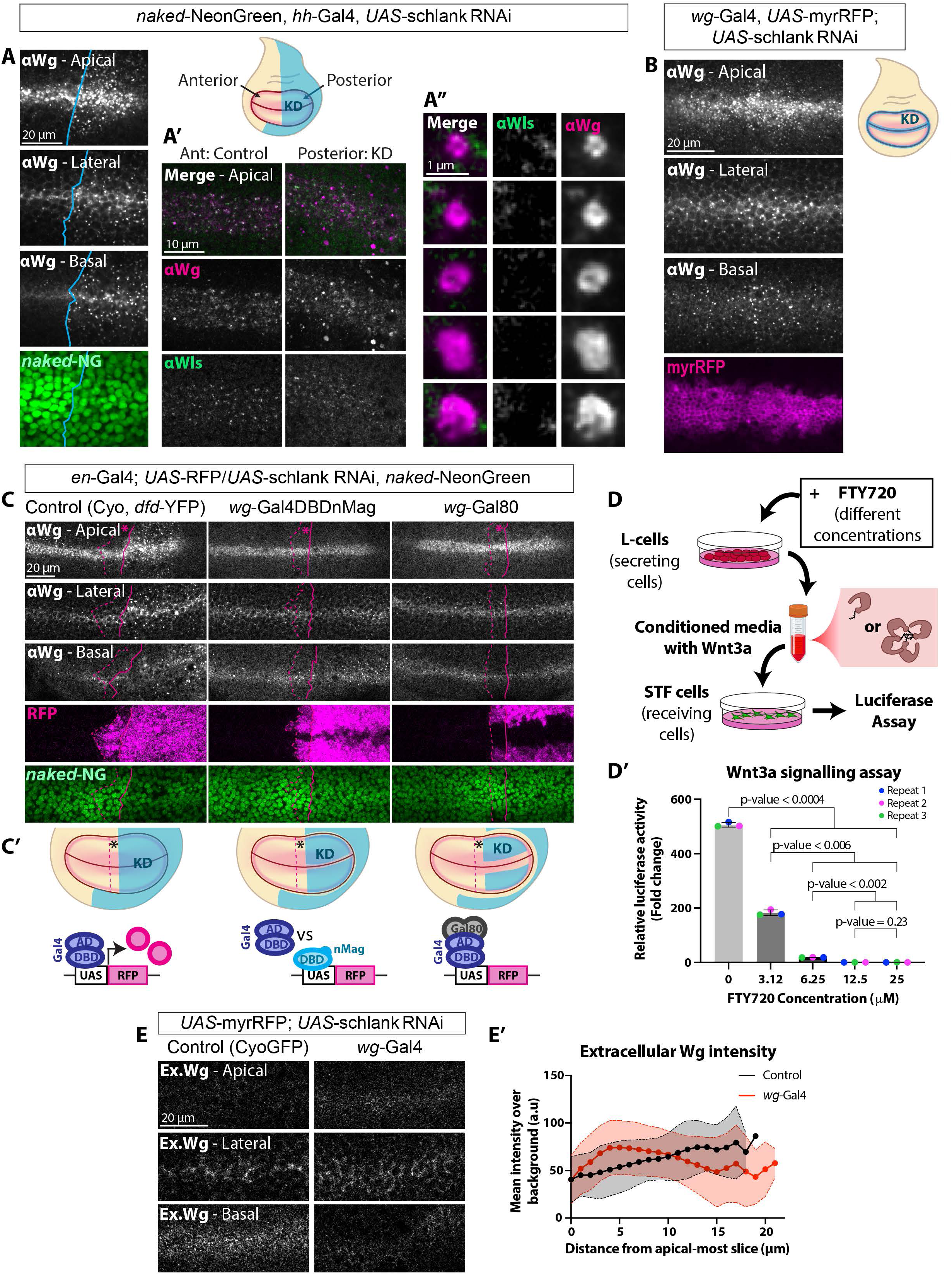
Reduced ceramide synthesis in Wg-producing cells causes Wg aggregation and apical release. **(A)** Total Wg and Wls immunoreactivity and GFP fluorescence from *nkd-NeonGreen* after *schlank* knockdown with *hh-Gal4*. Conventional confocal micrographs are shown in *A* and super-resolution in *A’-A’’. A’’* shows examples of WgLVs and large lumen-less punctae. The anterior-posterior boundary was recognised by anti-Ci staining (*blue* line). **(B)** Total Wg distribution after *schlank* knockdown with *wg-Gal4*. Gal4 activity is reported by myr-RFP. **(C)** Total Wg distribution after *schlank* knockdown in receiving cells. In preparations shown on the *left*, *schlank* RNAi is expressed throughout the posterior compartment with *en-Gal4*. In the other columns, *en-Gal4* is also present, but its activity is dampened in Wg-producing cells by Gal4DBDnMag (*middle*) or Gal80 (*right*), as outlined in *C’*. Asterisks mark a late domain of *en-Gal4* activity, recognised by faint *UAS-RFP* signal in the micrographs. Only incomplete *schlank* knockdown is achieved in this domain (see also FigS4D). **(D)** Signalling activity of Wnt3a-secreting L-cells treated with different concentrations of the ceramide synthase inhibitor FTY720. A summary of the assay is shown in *D*, and the results are shown in *D’*. Bars indicate mean luciferase activity from SuperTopFlash for each condition. N = 3 independent replicates, with each symbol being the average of three technical replicates. Statistical test: one-way ANOVA with Welch correction and Dunnett’s T3 multiple comparisons test. Error bars indicate SD. **(E)** Extracellular Wg distribution after *schlank* knockdown with *wg-Gal4*, with quantification along the apical-basal axis. Symbols indicate average signal along the apical-basal axis for all the wing discs analysed (N = 10 control and 11 knockdown discs), shading indicates SD.

To determine if *schlank* knockdown affects primarily secreting cells, receiving cells, or both, we developed means of knocking down *schlank* specifically in either cell population. Knocking down *schlank* in Wg-producing cells (using a *wg-Gal4* driver) suffices to trigger the formation of abnormal punctae both apically and basally (Fig5B), consistently with a role in producing cells. However, during early development, *wg-Gal4* is expressed throughout the wing primordium at low level and only refines later into a thin stripe (Alexandre et al., 2014; Couso et al., 1993; Ng et al., 1996; Phillips and Whittle, 1993; Williams et al., 1993). Therefore, one cannot exclude a possible effect of early knockdown in Wg-receiving cells. To overcome this limitation, we introduced *hh-Gal80*, which, as shown with a *UAS-RFP* transgene, suppresses Gal4 activity throughout the posterior compartment, except in a thin stripe where *wg-Gal4* is most expressed, ensuring knockdown only in Wg-secreting cells of the posterior compartment (FigS4C). Abnormal Wg punctae were seen in the posterior compartment, where *UAS-schlank^RNAi^* is expressed in a narrow stripe, as well as in the anterior compartment, where Gal80 is not expressed (FigS4C), confirming that *schlank* knockdown specifically in Wg-secreting cells leads to punctae formation. For the converse experiment, UAS-*schlank^RNAi^* was activated in the whole posterior compartment except in Wg-expressing cells (Fig5C). This was achieved by combining *en-Gal4* with either of two Gal4 inhibitors expressed from the endogenous *wg* promoter: Gal80 or Gal4-DBD-nMag. While Gal80 neutralizes the Gal4 activation domain (Ma and Ptashne, 1987), Gal4-DBD-nMag, one half of ShineGal4 (di Pietro et al., 2021), is expected only to weaken Gal4 activity by competing for Gal4 DNA binding sites. Indeed, expression of a *UAS-RFP* transgene shows that *wg-Gal80* represses Gal4 over a wider domain than *wg-Gal4-DBD-nMag* (Fig5C). In *en-Gal4, wg-Gal80, UAS-schlank^RNAi^*, no Wg punctae were apparent and signalling activity was normal. With *wg-Gal4-DBD-nMag*, which interferes with *en-Gal4* less well, Gal4 activity (and hence *schlank* knockdown) encroached the *wg* expression domain (as seen with the narrow band of RFP repression) and a mild phenotype became apparent: a few Wg punctae appeared at the basal side of the epithelium, while none could be detected in the apical and midlateral regions. In additional experiment, we found that, under condition of *schlank* depletion, optogenetically-induced newly produced GFP-Wg also formed punctae (FigS5), excluding the possibility that punctae form over time because of a traffic jam in the secretory pathway. These results show that, upon *schlank* depletion, Wg-producing cells release Wg in abnormal punctae.

In Drosophila imaginal discs, punctae formation correlates with reduced signalling, suggesting that Wg punctae could represent inactive Wg. To further evaluate this suggestion, we assessed the effect of reducing ceramide synthesis on the ability of mammalian cells to produce active Wnt. L-cells expressing Wnt3a were treated with FTY72, an general inhibitor of ceramide synthases (Berdyshev et al., 2009; Lahiri et al., 2009). After a week of treatment, conditioned medium was collected and its signalling activity was measured with a SuperTopFlash (STF) assay in HEK293 cells. FTY72 inhibited the conditioned medium’s signalling activity in a dose-dependent manner (Fig5D) in accordance with the effect of *schlank* depletion in Drosophila wing discs.

So far, our results show that both endocytosis blockade or *schlank* depletion leads to the formation of abnormal Wg punctae. One difference between the two conditions is that, with endocytosis blockade, punctae are confined to the apical surface, whereas with *schlank* depletion, they are found throughout the apical-basal axis. The effect of *schlank* RNAi cannot be attributed to inhibition of endocytosis since endocytosis can proceed in the absence of Schlank (Hull et al., 2024; Pepperl et al., 2013) (see Discussion). Therefore, the *schlank* RNAi-induced punctae could form at the apical surface before being endocytosed and/or along the apical-basal axis during transcytosis. Irrespectively, it appears that *schlank* depletion disrupts Wg trafficking since, upon *schlank* knockdown, the basal extracellular Wg gradient is lost and the level of apical and subapical extracellular Wg protein increases (Fig5E).

### Dlp ‘solubilizes’ Wg punctae

The localisation of Wg at the inner periphery of WgLVs indicates membrane association. In contrast, the Wg punctae that form upon *schlank* knockdown lack a visible lumen (Fig5A), suggesting that they could represent insoluble aggregates. Wnts can be maintained in solution by lipid binding proteins (reviewed in (Alvarez-Rodrigo et al., 2023). For example, in vitro, Wnt proteins can transfer from Wls to sFRP2, WIF1 or Afamin albeit only in the presence of detergent (de Almeida Magalhaes et al., 2024). These proteins are not present in Drosophila but the Wg lipid can also be shielded *in vivo* by a hydrophobic tunnel in the glypican Dlp (McGough et al., 2020), which is present in endosomes in wing imaginal discs (Gallet et al., 2008). Super-resolution microscopy of discs endogenously expressing SNAP-Wg labelled with SNAP-Cell TMR-Star shows that Dlp localises to some, but not all, WgLVs (Fig6A). Therefore, although Wg could bind Dlp in the endocytic pathway, this interaction is not essential for Wg’s association with the inner face of endocytic vesicles. We also found that WgLVs form in the absence of Dlp (*dlp[KO])* with their size, number and amount of Wg within the wild-type range (Fig6B-C), confirming that Dlp is not required for membrane association of Wg in endosomes. Nevertheless, punctae do appear at the basal surface of *dlp[KO]* tissue (Fig6B), suggesting an essential function there. We conclude that Dlp is not required to host the Wg lipid at the inner surface of WgLVs but may help prevent aggregation at the basal surface before onward transport to receiving cells.

**Figure 6.**
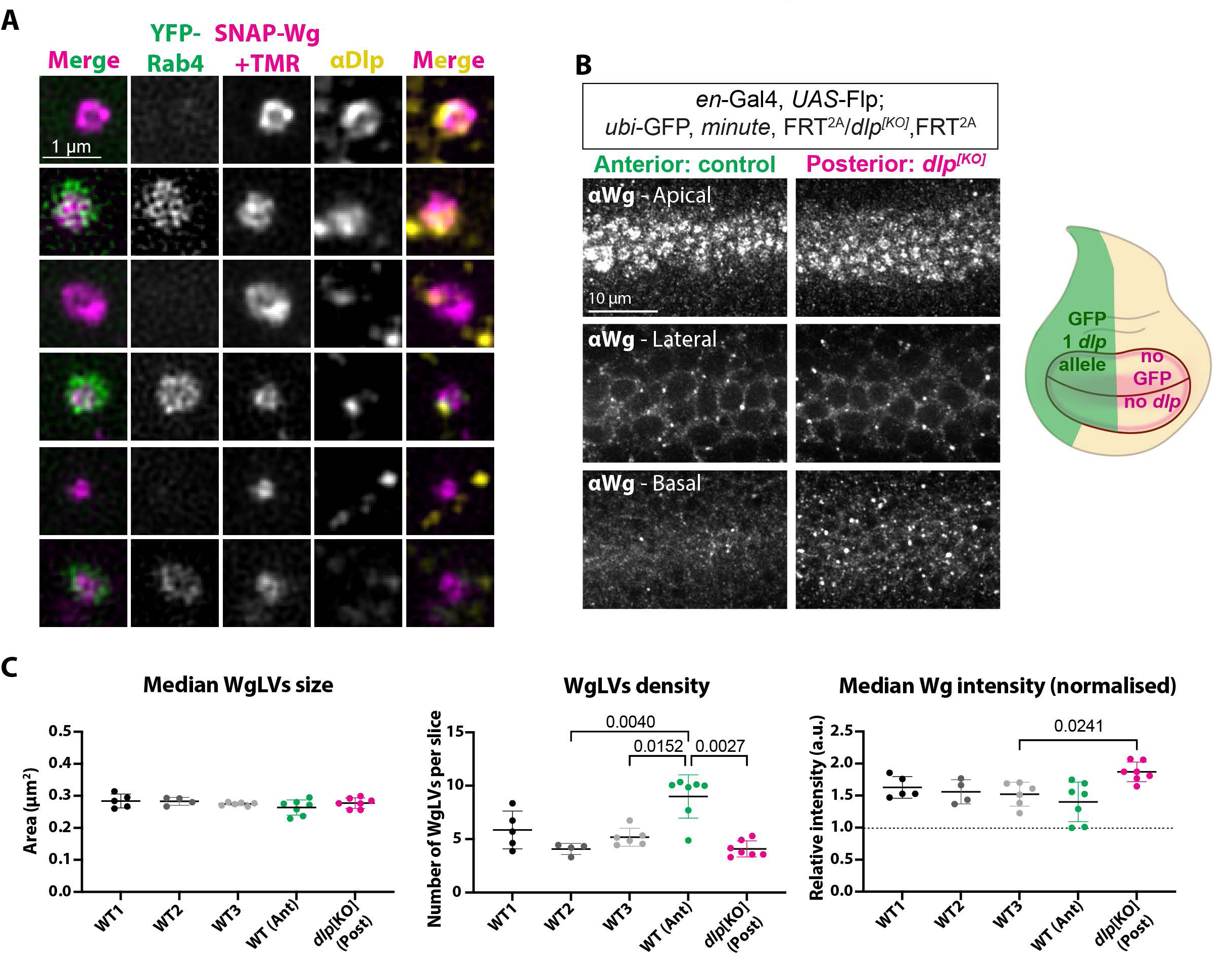
Dlp is required for basolateral Wg secretion but not for localization to WgLVs. **(A)** WgLVs in wing discs expressing YFP-Rab4 and SNAP-Wg expressed from the corresponding endogenous locus and stained with anti-Dlp (SNAP-Wg was used to detect Wg because anti-Wg and anti-Dlp are both from mice). Some WgLVs, but not all, contain Dlp irrespectively of the presence of Rab4. **(B)** Total Wg in discs lacking *dlp* (generated by mitotic recombination) in the posterior compartment. The heterozygous anterior compartment serves as a control. **(C)** Quantification of WgLV parameters from the data exemplified in *(B)*. For each disc, we calculated a median WgLV size, WgLV density and normalised median Wg intensity. Three independent cohorts of wild-type wing discs are also shown, in *grey*. From left to right, N = 5, 4, 6, 7 and 7 discs, with >80 WgLVs analysed per disc. Statistical test: one-way ANOVA with Welch correction and Dunnett’s T3 multiple comparisons test. Graph bars: mean with SD.

There is evidence that the Wnt lipid can associate directly with the lipid bilayer since purified Wnt3a can be packaged in liposomes (Janeczek et al., 2017; Morrell et al., 2008; Tüysüz et al., 2017). We confirmed that purified GFP-Wg can be incorporated in 1,2-Dimyristoyl-*sn*-Glycero-3-Phosphocholine (DMPC) liposomes, mostly at the outer surface, as inferred from a trypsin digest assay (Fig7A-B). Most of the Wg-associated liposomes that we generated were too small for visualisation of a lumen (median GFP+ particle area = 0.018 μm^2^). However, the few large liposomes (GFP positive particle area > 0.03 μm^2^) that could be identified manually were found by optical pixel reassignment and deconvolution to have a recognisable lumen (Fig7C). In control experiments, we assessed the behaviour of purified GFP-Wg diluted in buffer with or without CHAPS detergent (Fig7D-E). While punctae formed only rarely in the presence of detergent, many appeared in the preparation lacking CHAPS, suggesting that they represent aggregates that form because the Wg lipid is not shielded. After 1 h incubation at room temperature without CHAPS, punctae number decreased and their individual fluorescence intensity increased, with total fluorescence intensity remaining approximately constant, suggesting that the aggregates may be fusing over time. It is tempting to suggest that these structures mimic the abnormal Wg punctae that form in vivo upon *schlank* knockdown.

**Figure 7.**
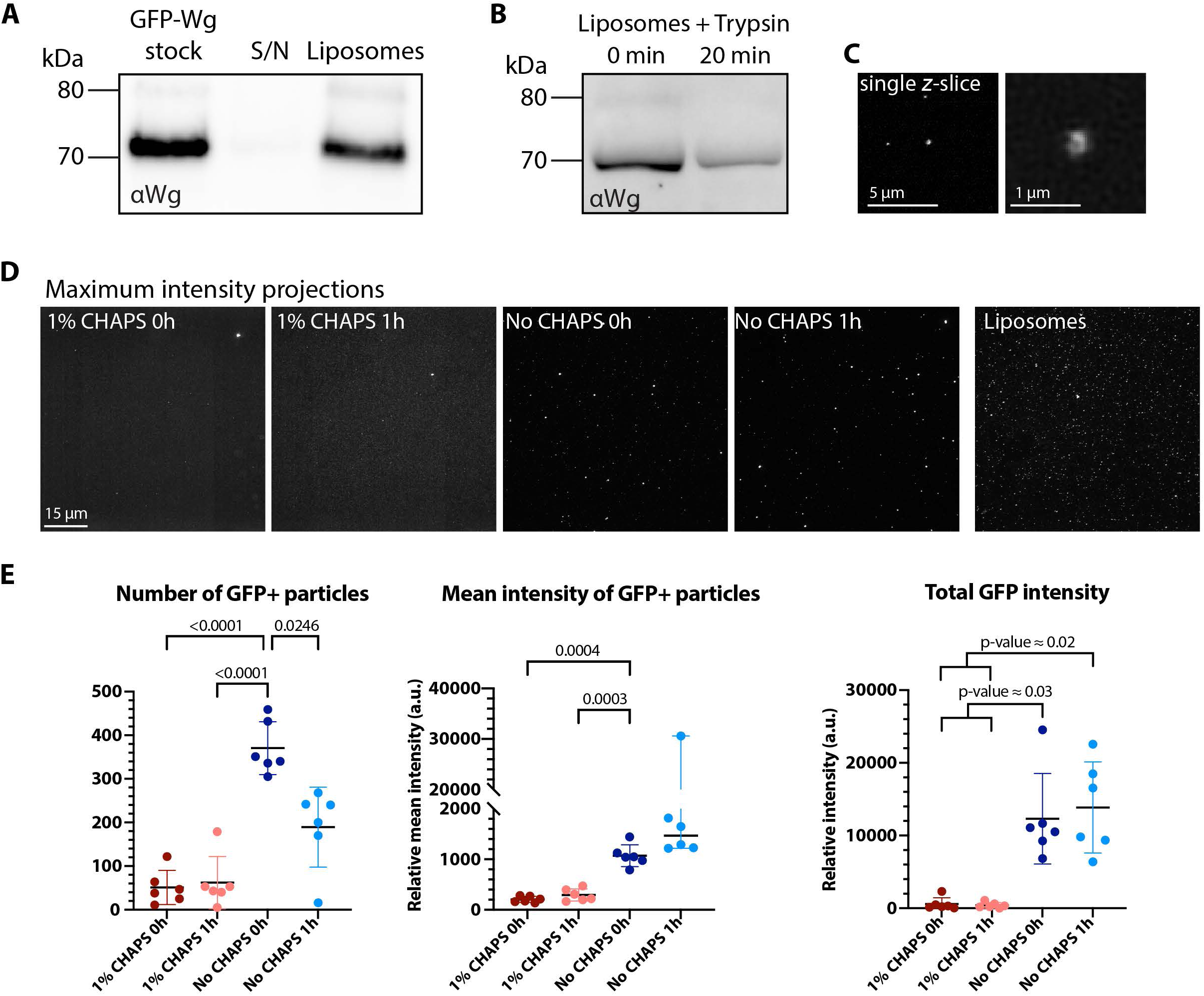
Liposome packaging prevents GFP-Wg aggregation in vitro. **(A)** Western blot (representative of three repeats) showing the purified GFP-Wg stock used for incorporation in liposomes (*lane 1*), along with the supernatant (*S/N, lane 2*) and pellet (*liposomes, lane 3*) obtained after centrifugation of the liposome preparation. **(B)** Western blot (representative of three repeats) of GFP-Wg liposomes treated with trypsin, either immediately quenched (0 min) or allowed to act for 20 min at 37°C. **(C)** Super-resolution (single optical section) image of a GFP-Wg liposome. In this example, the liposome was large enough for a lumen to be resolved. **(D)** Super-resolution images (shown as maximum intensity projections) of purified GFP-Wg diluted with or without CHAPS and incubated at RT for 0 or 1 h. Since background signal is much higher in the detergent sample than the non-detergent sample (perhaps because of GFP-Wg in solution) brightness/contrast levels cannot be compared. A super-resolution image of a liposome preparation is shown for comparison. **(E)** Parameters from the data exemplified in *(D)*: number and mean intensity of the GFP-Wg aggregates, and total GFP intensity of all aggregates in the images. N = 6 different areas of the petri dish per condition. Statistical test: one-way ANOVA with Welch correction and Dunnett’s T3 multiple comparisons test. Graph bars: mean with SD.

The above in vitro data suggest that ceramide synthase depletion could trigger Wg aggregation by overcoming the bilayer’s capacity to accommodate the Wg lipid. If punctae do represent aggregates, the removal of Dlp, which can provide an additional landing platform for the Wg lipid, would be expected to exacerbate the formation of *schlank* RNAi-induced punctae. We therefore tested whether partial loss of Dlp had an effect on the *schlank* RNAi phenotype. Inactivation of *FRT-dlp-FRT* with *vg-Gal4 UAS-Flp,* has no overt discernible phenotype on Wg trafficking, perhaps due to perduring Dlp activity (*vg-Gal4* is activated later than *en-Gal4*). Yet, it led to a marked enhancement of the abnormal punctae in a *schlank* RNAi background (Fig8A), especially at the basal surface. Therefore, Dlp appears to limit the formation of Wg aggregates. Moreover, Dlp protein still localises to residual WgLVs in *schlank*-deficient cells (Fig8B), an indication that *schlank* RNAi does not impede Dlp recruitment to WgLVs.

**Figure 8.**
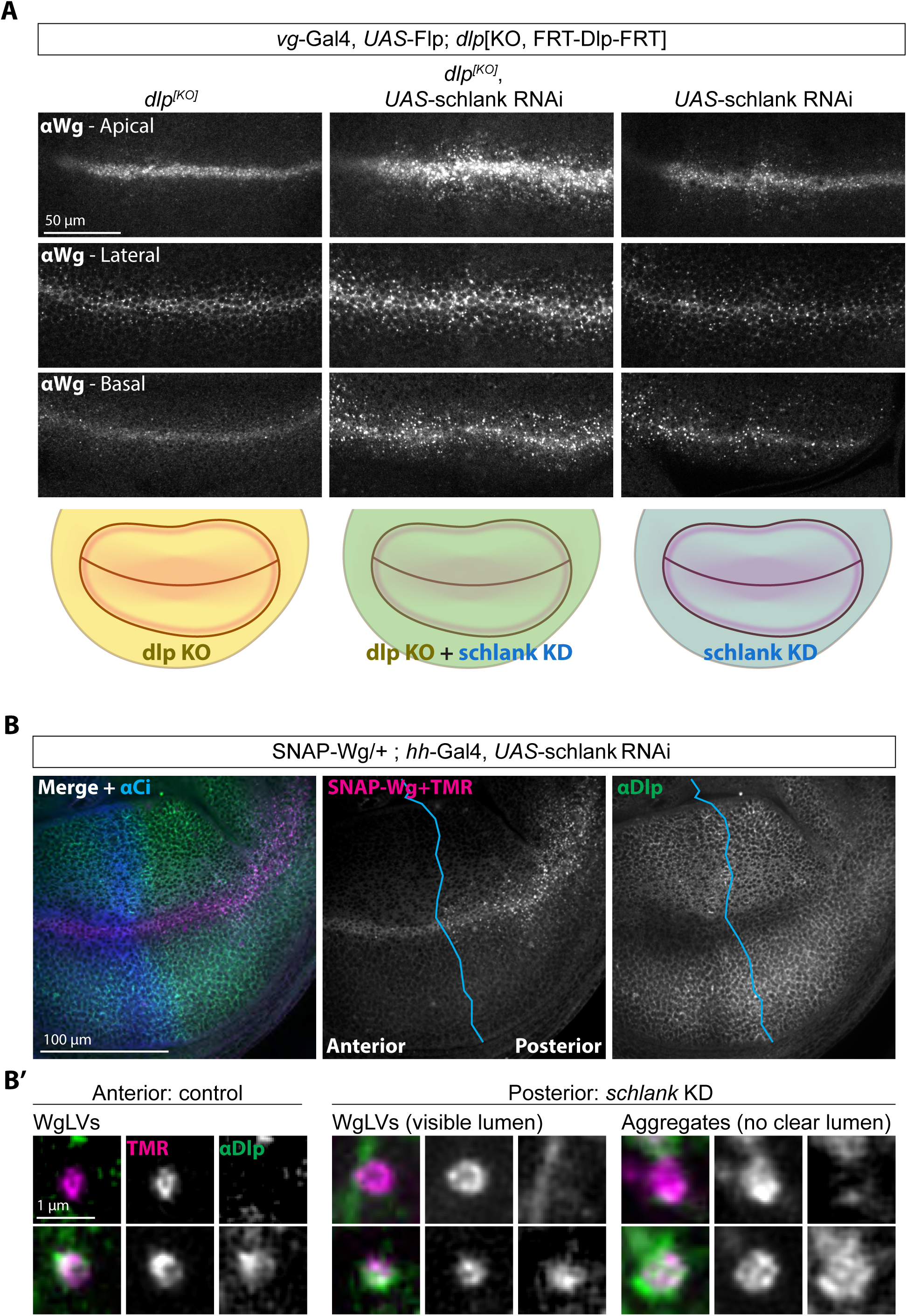
Synergistic effect of reducing Schlank and Dlp activity. (A) Total Wg distribution after *dlp* inactivation (*left), schlank* knockdown (*right*) or both (*middle*) in the *vg-Gal4* domain. (B) Total Wg (from SNAP-Wg labelled with SNAP-TMR) and Dlp (from immunofluorescence) after *schlank* knockdown under the control of *hh-Gal4*. Anti-Ci marks the anterior compartment. Both WgLVs and Wg punctae appear in the *schlank* knockdown compartment while only WgLVs are seen in the control compartment. Dlp was detectable in some but not all WgLVs and in some but not all the aggregates.

Having shown that loss of Dlp enhances the phenotype of ceramide synthase depletion, we set out to ask if Dlp overexpression would alleviate it. Indeed, overexpression of wild-type Dlp suppressed the formation of Wg punctae (Fig9), though not the signalling defect caused by *schlank* RNAi (see Discussion). By contrast, overexpression of a form of Dlp that was mutated to prevent opening of the lipid-binding tunnel (Dlp^MN/CC^, “Clamp”) was inefficient at preventing punctae formation, highlighting the importance of lipid shielding. Since Dlp Clamp did partially alleviate the phenotype of *schlank* RNAi, it is possible that GAG chains on Dlp Clamp could contribute to punctae suppression. We conclude that depletion of ceramide synthase disrupts Wg localisation to the inner leaflet of endosomes, leading to the formation of aggregates that can be solubilised by the lipid-binding tunnel of Dlp.

**Figure 9.**
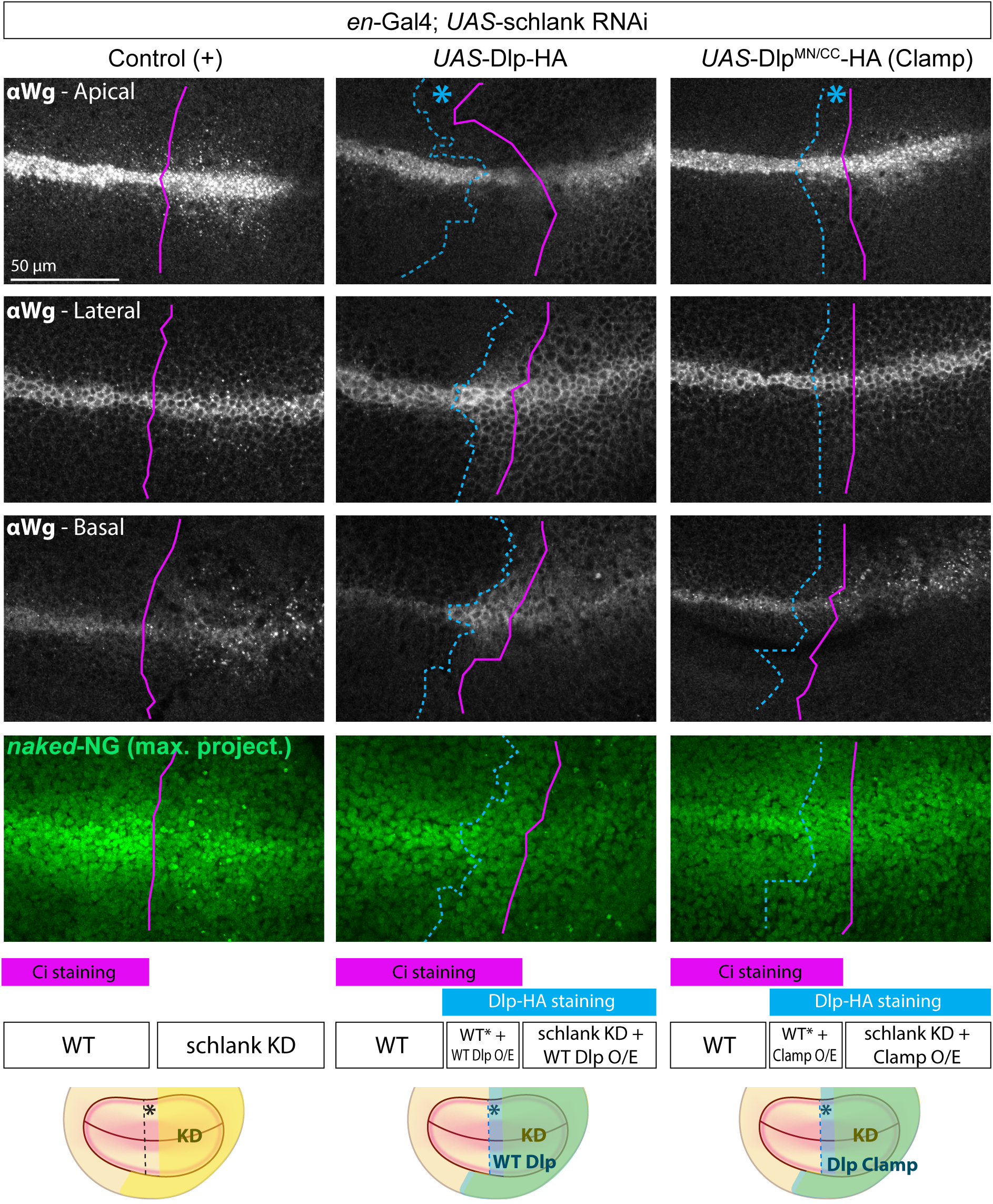
Lipid-shielding activity of Dlp alleviates schlank RNAi-induced Wg aggregation. Total Wg (single *z-*slices) and *naked-NeonGreen* (maximum intensity projections) in discs expressing *schlank* RNAi alone, or in combination with Dlp-HA or Dlp^MMCC^-HA (no lipid binding activity), all under the control of *en-Gal4.* Magenta line shows compartment boundary, recognised by anti-Ci staining. In the area marked with an asterisk, late activation of *en-gal4* is sufficient to trigger Dlp expression but not to appreciably knock down *schlank*. Therefore, three conditions are created in the discs shown in the *middle* and *right* columns: wild type, near wild type *schlank* activity with overexpressed Dlp variants, and *schlank* knockdown with overexpressed Dlp variants.

## Discussion

Wnts are unique among secreted proteins in that they are post-translationally appended with a palmitoleoyl moiety (Resh, 2013). This lipid, which is essential for signalling activity, must be shielded for progression through the secretory pathway. This is achieved by a Wnt-specific chaperone, Wls, which escorts Wnts from the ER to the plasma membrane, keeping the Wnt lipid in a hydrophobic tunnel (Qi et al., 2023). For active Wnt to be released, the Wnt lipid must disengage from this tunnel, perhaps through a conformation change in Wls. The trigger of dissociation and the immediate destination of the Wnt lipid after dissociation have not been elucidated so far. We have approached this question by first investigating where, at the subcellular level, Wg (the archetypical Drosophila Wnt) dissociates from Wls in wing imaginal discs (Fig10). Upon reaching the plasma membrane, Wls is rapidly endocytosed by virtue of an AP2 binding site on a intracellular loop, before being recycled to the ER/Golgi (Belenkaya et al., 2008; Gasnereau et al., 2011; Pan et al., 2008; Yang et al., 2008; Yu et al., 2014). We find that Wg is also re-internalised. This may appear counterintuitive for a protein that needs to be secreted, but this step is necessary for eventual release in the extracellular basolateral space (Yamazaki et al., 2016), where the signalling gradient is generated. We found that Wg internalisation is rapid, at least after release from endocytosis inhibition (Fig3). The two proteins must part ways in the endocytic pathway since Wg subsequently traffics, without Wls, to the inner membrane of specialised endosomes that we name WgLVs.

**Figure 10.**
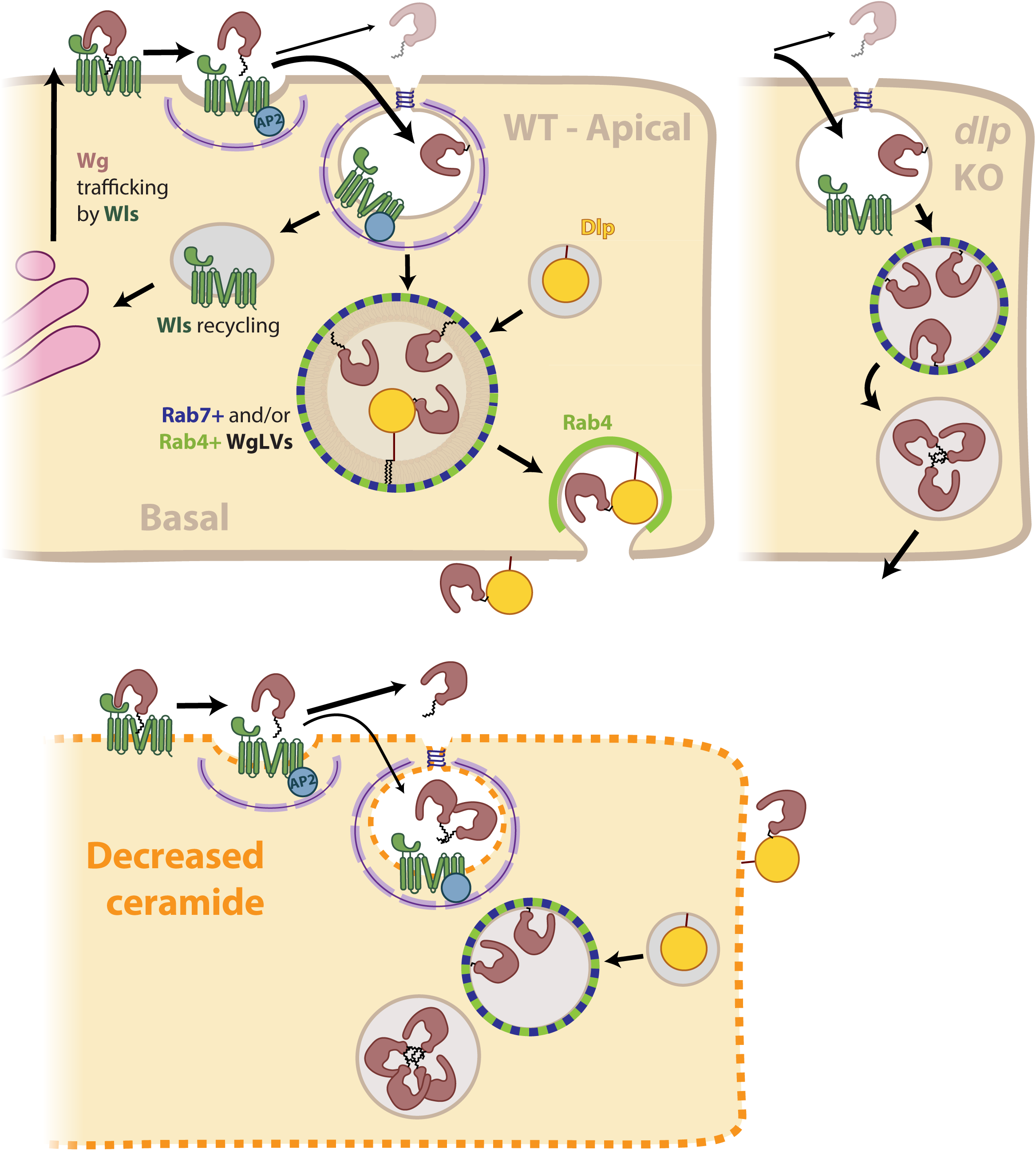
Appropriate membrane lipid composition and endocytosis ensure that Wingless does not aggregate in secreting cells after Wls dissociation. Diagrammatic view of Wg trafficking in wild type, *schlank*-deficient, or *dlp*-deficient Wg-secreting cells. In the wild type, Wg-Wls dissociation commences at the apical plasma membrane rapid but endocytosis and incorporation in the lipid bilayer prevents apical release. Wg accumulates into WgLVs marked by Rab7 and/or Rab4, where Dlp is progressively recruited until basolateral Wg release. In the absence of Dlp, Wg still reaches WgLVs but aggregates over time in the basal region. Upon ceramide synthesis knock down, orderly transfer from Wls to the bilayer is disturbed, resulting in abnormal apical Wg release and accumulation of Wg aggregates.

We found WgLVs to be coated with Rab4, Rab7, or both. Since Rab4 is a marker of recycling endosomes, it is tempting to consider that Rab4-positive WgLVs could be a transit station towards the basolateral plasma membrane for release. Although colocalization of Rab4 and Rab 7 may be unexpected, it is not unheard of. Rab7 has been found to overlap with other markers such Rab4, Rab11 or Rab19 (Fan et al., 2020; Prince et al., 2019; Yu et al., 2024). Perhaps double positive Rab7/Rab4 WgLVs represent that progress from Rab7- to Rab4-positive endosomes. However, we cannot exclude the possibility that some Rab7-positive WgLVs could be carrying excess Wg towards lysosomes for degradation. Although we expect Wg to transit through Rab5-positive endosome shortly after endocytosis, no Rab5-positive WgLVs were recognised, perhaps because early endosomes were too small for a lumen to be resolved with our super-resolution microscopy pipeline (FigS3A).

Acute inactivation of Dynamin leads to the formation, at the apical surface, of abnormal Wg punctae devoid of Wls, indicating that dissociation can take place there. However, this must occur at a slow rate since a substantial amount of Wg remains associated with the apical surface. In this experimental set up, release could possibly occur from the plasma membrane or endocytic pits, which still form in the absence of dynamin activity (Koenig and Ikeda, 1989). However, punctae also form after inhibition of Clathrin, which is required for coated pits to form. We conclude therefore that Wg release can occur from the apical plasma membrane, before any increase in membrane curvature or acidification, two features that have been considered as possible triggers of release (Alvarez-Rodrigo et al., 2023; Coombs et al., 2010; Sharma and Chaudhary, 2024). Although we cannot exclude that these processes could speed up Wg-Wls dissociation in early endosome, we suggest that the initial trigger must be a distinct feature of the plasma membrane, such as for example specific lipid composition or flattening of the lipid bilayer.

Whatever its trigger, Wg-Wls dissociation is likely to be sensitive to slight changes in Wls’ conformation, as suggested by Wls tagging efforts. Specifically, insertion of GFP in the fourth luminal loop (*wls[ExGFP]*) has seemingly no adverse effect since flies expressing this form from the endogenous locus are homozygous viable (Yu et al., 2020). Yet this modification dominantly causes mild Wg retention within Wg-producing cells (FigS2). Unlike wild-type Wls, Wls[ExGFP] can be found in WgLVs, suggesting that the GFP moiety may hamper Wg release from Wls. This observation may explain why Wls and Wg have been suggested to separate only in maturing acidic endosomes (Sharma and Chaudhary, 2024). It also suggests that a mild structural change can affect the ability of Wls to release Wg. Therefore, release could conceivably be normally triggered by a conformational change induced by the plasma membrane’s local environment.

Recent structural analysis of Wls showed that its central cavity can accommodate a phospholipid from the bilayer, and that its presence could promote a stable Wls-Wnt confirmation (Nygaard et al., 2021; Qi et al., 2023; Zhong et al., 2021). In light of this observation, we revisited a report that ceramide depletion caused by *schlank* RNAi impairs Wg trafficking and signalling. We found that *schlank* knockdown, specifically in secreting cells, leads to the formation of Wg punctae, which may represent inactive insoluble aggregates. Note that, while we cannot exclude a role of ceramides in receiving cells, e.g. for signalling activity (Pepperl et al., 2013), our analysis demonstrates the role of ceramide synthesis in Wg-secreting cells. Depletion of *schlank* is not thought to hinder endocytosis, since it does not affect Dextran uptake (Pepperl et al., 2013) or cause the gas-filling defects characteristic of impaired endocytosis during tracheal development (Hull et al., 2024). It is unlikely therefore that the formation of Wg punctae in *schlank*-deficient Wg secreting cells is due to impaired endocytosis.

How does reduction of ceramide synthesis affect Wg secretion? One possibility, suggested by the structure of the Wnt-Wls complex (Qi et al., 2023), is that altered lipid composition lowers occupancy of Wls’ central cavity by a specific lipid, e.g. a sphingolipid, weakening the affinity of Wls for Wg (the premature release hypothesis). Another possibility is that altered lipid composition of the bilayer reduces its ability to accommodate the Wg lipid (the impaired destination hypothesis). According to either scenario, upon ceramide synthase depletion, excess Wg would accumulate in the lumen of WgLVs and the Wg lipid would become exposed, triggering aggregation. Accordingly, overexpressed Dlp, which is known to shield the Wg lipid (McGough et al., 2020) was found to reduce punctae formation. Although we cannot exclude the possibility that another feature of the apical plasma membrane (e.g. flattening of the bilayer) could trigger Wg-Wls dissociation, our results highlight the importance of lipid composition. Future in vitro work will be required to test how different lipid preparations affect Wnt packaging into liposomes and/or Wnt-Wls complex stability.

## Acknowledgements

We would like to thank Konrad Basler for kindly providing us with anti-Wls antibody and Drosophila strains that were key to our project. We are grateful to Joachim Kurth from the Crick’s Fly Facility for performing the Drosophila injections and the rest of the Fly Facility team for further support. For their advice and support with microscopy and image analysis we thank Rocco D’Antuono, Matt Renshaw, Donald Bell, Dave Barry and Todd Fallesen (Advanced Light Microscopy Facility at the Francis Crick Institute); and Alan Wainman (Sir William Dunn School of Pathology, University of Oxford). We are also grateful to current and past members of the Vincent laboratory, the Crick Fly community, our “Pass the Wnt” collaborators (Clausen and Salbreux laboratories), Jeremy Carlton, Clive Wilson, and the WntUK community for valuable discussions and constructive feedback. Special thanks to Ian McGough for his insight and support throughout this project. This work was funded by the Wellcome Trust (collaborative award 223133/Z/21/Z to E.Y.J and J.-P.V. and discovery award 214259/Z/18/ to P.B.), by SFB 645 (to R.B.), Institut Curie, CNRS, INSERM and ANR (to Y.B.) and core funding from the Francis Crick Institute to J.-P.V. (FC001204). The Francis Crick Institute receives its core funding from Cancer Research UK, the UK Medical Research Council, and the Wellcome Trust.

## Materials and Methods

### Drosophila melanogaster *stocks and husbandry*

Fly strains were raised on standard agar media at 25°C, unless stated otherwise. Standard fly handling techniques were employed (Roberts, 1998). DNA injection was performed by the Francis Crick Institute fly facility unless otherwise indicated. *w1118* was used as wildtype, for all other fly lines used see Supplementary Table 1.

For live imaging of GFP-Wg embryos, embryos were collected on cranberry–raspberry juice plates (25% cranberry–raspberry juice, 2% sucrose and 1.8% agar) supplemented with fresh yeast. Embryos were collected for 1.5 h at 25°C, aged for 1 h, mounted, aged at 25°C until cellularisation (approx. 3h) and monitored on the microscope until they reached stage 9 - 11 (4 – 6 h elapsed time). For mounting, embryos were dechorionated on double-sided tape and mounted on a strip of glue onto a 35 mm glass bottom Petri dish with a 14 mm micro-well (MatTek). After desiccation for 1 min at 25°C, embryos were covered in Voltalef oil (ARKEMA).

### Generation of transgenic flies

A strain expressing *SNAP-Wg* from the *wg* locus was generated by reintegrating SNAP-Wg-RIV{pax-Cherry} into *wg[KO;attP]* line (Alexandre et al., 2014). To build SNAP-Wg-RIV{pax-Cherry}, Wg-encoding sequences were obtained from a RIV{pax-Cherry} variant harbouring full-length *TurboID-Wg* cDNA along with 135 bp of 5′ untranslated region (UTR) and 1,200 bp of 3′ UTR (Androniciuc et al., 2024). NEBuilder HiFi DNA Assembly (NEB) was used to replace the TurboID sequence with a SNAP-tag sequence obtained from plasmid pSNAPf-N1, a gift from Michael Davidson (Addgene, plasmid 58187), inserted between the Wg signal peptide and a GGGNGRG linker (see primers in Supplementary Table 2). The *SNAP-Wg* strain was homozygous viable and presented no defects in wing shape or margin bristles.

To generate a strain expressing GFP-Chc from the *chc* locus, we used the method described in (Poernbacher et al., 2019), starting from the plasmid IOGFP@NTV-Pax-Cherry. Briefly, the CRISPR site and homology arms were chosen so that after the final recombination event the first codon of GFP would correspond to the translation start of the endogenous *chc* locus (see Supplementary Table 2). Gibson assembly was used to insert the homology arms into the IOGFP@NTV-Pax-Cherry plasmid pre-digested with BsaI and SapI, and the gRNAs into a pCFD4 vector. Both vectors were co-injected into a Cas9-expressing strain (*nos-Cas9-mSA*). Successful candidates (females, as the males were not homozygous viable) were identified by expression of GFP and *pax-mCherry*, which was then excised via Cre-mediated recombination. The resulting strain, which expresses N-terminally tagged Chc from the endogenous *chc,* with a linker comprising the residual LoxP site and additional 11 G/S repeats (aminoacid sequence at the GFP-Chc fusion border after Cre-mediated excision: […]MDELYK/ITSYNVCYTKLC/(GS)x11/TQPLPIRFQEHLQ[…]).

The *UAS-OptoCD8trap* line was generated by introducing the 2xnMagHigh1 and 2xpMagHigh1 modules (di Pietro et al., 2021) separated by a T2A sequence, into a *UAS-CD8NB* plasmid, which comprises sequences encoding amino-acids 1-227 of the mouse CD8 sequence (NP_001074579), an 8 amino-acid Gly/Ser linker (including a unique Xho site) and an anti-GFP VhhGFP4 nanobody (Saerens et al., 2005). The *UAS-CD8NB* plasmid was linearised with Xho and the nMag2ApMag ORF was then inserted between the CD8 and nanobody encoded sequences (see primers in Supplementary Table 2).

The *ubi-Cry2olig-Vhh-mCherry* line was generated by combining sequences encoding Cry2olig (López-Gay et al., 2020) and anti-GFP Vhh nanobody (Caussinus et al., 2012) and an intervening six amino-acids Gly/Ser linker into the *ubi-Mod-Poly* vector (David et al., 2005). The F-Ubi-Cry2 and R-Vhh-Cry2 primers were used to amplify the Cryolig sequence and the F-Cry2-Vhh and R-Ubi-Vhh primers were used to amplify the Vhh sequence and the linker (see primers in Supplementary Table 2). The Cry2olig-Vhh product was then cloned at the BglII and XbaI sites of the Ubi-Mod-Poly vector.

The *wg-Gal80* line was generated by inserting the RIV^Gal80^ plasmid (Baena-Lopez et al., 2013) in *wg[KO;attP]* flies (Alexandre et al., 2014) using φC31 site-directed integration. RIV^Gal80^ was integrated the attP site of *hh[KO;attP]* flies (Baena-Lopez et al., 2013) to generate the *hh-Gal80* strain.

To generate the *FRT-schlank-HA-FRT* strain, a 4 kb genomic DNA (gDNA) encompassing exons 2-6 and the *schlank* 3’ UTR was digested with NotI and AscI and cloned into the NotI and Asc I sites of RIVFRT MCS2FRT QF (Baena-Lopez et al., 2013). This construct was then integrated by BestGene Inc. (Chino Hills, CA, USA) into *schlank[KO;attP]*’s attP site, which is located in the non-coding first intron (Sociale et al., 2018). *FRT-schlank-HA-FR*T flies were phenotypically indistinguishable from wild-type flies (*w^1118^*).

To generate *UAS-GFP-Wg* (fully functioning GFP-Wg fusion), the 10x UAS pJFRC81 vector (PMID: 22493255) was digested with AgeI and NotI for insertion GFP-Wg encoding DNA and an upstream FRT-STOP-FRT cassette (GFP immediately Wg downstream of the signal peptide and followed by the linker IASKPKGASVRA). The resulting construct was integrated at attP1 by ΦC31-mediated integration and the stop cassette was excised in candidate transgenic flies before establishment of the final strain.

### Acute endocytosis block

To inhibit Dynamin temporarily, *shibire^ts^* wandering L3 larvae were treated as in (Yamazaki et al., 2016). For acute inhibition of clathrin, the crosses *GFP-chc//; wg-Gal4/CyoTbRFP* females x *UAS-OptoCD8trap /TM6B* males, or *GFP-chc//* females *x ubi-Cry2oligVHH-Cherry//* males were kept in the dark (vials wrapped in aluminium foil inside a cardboard box). Vials were flipped and wandering 3^rd^ instar males were selected under illumination covered with three layers of Deep Straw colour filter LP015 (HT-range, Chris James Lighting Filters). Control larvae were dissected in the same set up and fixed immediately. Test larvae were placed in PBS in a Petri dish and illuminated for 10 min using a BlueCell device equipped with an array of 460 nm LED strips (Gehrig et al., 2017) before being transferred to a petri dish with wheat food in a 25°C room with constant light, for a total 30 min or 1.5 h light exposure (see Fig4) before fixation.

### Immunostaining

For ‘total’ immunostaining (labelling both intra and extracellular protein) of wing discs, wandering 3^rd^ instar larvae were dissected in PBS and fixed for 30 min in 4% formaldehyde in PBS. Discs were washed thrice in PBS, permeabilised in 0.2% Triton-X100 in PBS for 10 min and blocked in 5% Bovine serum albumin (BSA) in PBT (PBS + 0.2% Triton-X100) for 10+ min. Samples were incubated in primary antibodies overnight at 4°C in 5% BSA-PBT, washed thrice in PBT, incubated in secondary antibodies in 5% BSA-PBT for 2 h at room temperature (RT), washed again thrice in PBT and mounted in SlowFade diamond (ThermoFisher, S36963), using a 1.5H high precision cover glass (Marienfield Superior, 0107222) and slides (Epredia, X1XER308B) sealed with CoverGrip (Biotium, 23005). Note that a different protocol was used for SNAP-Wg labelling (see next section).

Extracellular immunostaining was performed as previously described (Beckett et al., 2013). Control and test discs were dissected, fixed and stained together in the same tube to minimise technical variability. They were distinguished based on the presence or absence of Myr-RFP.

The following primary antibodies were used: mouse anti-Wingless (1:500, DSHB 4D4), rabbit anti-Wls (1:1000, generous gift from Konrad Basler, (Port et al., 2008)), goat anti-GMAP (1:1000, DSHB GMAP), rabbit anti-HA (1:800, CST #3724), rat anti-DE-cadherin (1:50, DSHB DCAD2), mouse anti-Rab7 (1:10, DSHB Rab7-s), rat anti-Ci (1:100, DSHB 2A1-s) and mouse anti-Dlp (1:50, DSHB 13G8).

Secondary antibodies used were anti-rabbit AlexaFluor488, anti-mouse Cy3, anti-rat AlexaFluor647 (1:400, Jackson ImmunoResearch); anti-rabbit CF405S, anti-rat CF405S (1:1000, Biotum); anti-rabbit AlexaFluor647, anti-mouse AlexaFluor405, anti-mouse AlexaFluor555, anti-mouse AlexaFluor647, anti-goat AlexaFluor647 (1:500, Invitrogen). We also used GFP-Booster AlexaFluor488 (1:500, Proteintech ChromoTek GB2AF488-50) and Hoechst 33342 (1:500, Invitrogen H3570).

### SNAP-labelling of wing discs

Wing discs of larvae expressing *SNAP-Wg* were dissected, fixed and washed as above. For labelling after permeabilization (FigS1Ci, Fig2D – surface649), wing discs were permeabilised with 0.5% Triton-X100 in PBS for 20 min, washed once in PBS, incubated in SNAP labelling reagent diluted to a final concentration of 2 μM in PBS for 15 min at RT while mixing, then washed once before proceeding with the rest of the immunostaining as normal. For labelling prior to permeabilization (FigS1Cii, Fig6A, Fig8B), wing discs were incubated in SNAP labelling reagent as above, washed once and permeabilised as normal in 0.2% PBT. SNAP labelling reagents used: SNAP-surface488 (NEB, S9124S), SNAPCell-TMR-Star (NEB, S9105S) and SNAP-surface649 (NEB, S9159S).

### GFP-Wg purification

The plasmids pExpresS2_Zeocin_GFP-Wg and pExpresS2_Puromycin_ Dlp^ΔGPI^-V5 were generated by subcloning pre-existing pMT plasmids (McGough et al., 2020) into pExpresS2 vectors digested with AgeI and NotI (ExpresS2ion Biotechnologies) using NEBuilder HiFi DNA Assembly (NEB) and the primers indicated in Supplementary Table 2. The GFP-Wg-expressing plasmid was then transfected into S2 parental cells (ExpresS2ion Biotechnologies) and selected over three weeks with Zeocin. Once the cell line was established, the Dlp plasmid was transfected so that secreted Dlp would stabilize secreted GFP-Wg in the supernatant, and selection occurred over two weeks using Puromycin. Cells were grown in EX-CELL^®^ 420 Serum-Free Medium (Sigma, 14420C) supplemented with Pen/Strep antibiotics and 10% FCS. The cells were tested for the presence of the two proteins in the supernatant by western blot, then expanded for purification. Cells were split twice a week, at 8x10^6^ cells/ml in EX-CELL^®^ 420 Serum-Free Medium, supplemented with Pen/Strep antibiotics.

To purify GFP-Wg, a 5ml HiTrap Heparin column (Cytiva, 17040701) was equilibrated with Buffer A (pH7.5, 25 mM HEPES, 50 mM NaCl). Cells were harvested at 3000 rpm for 15 min; the supernatant was filtered and loaded at 5 ml/min onto the HiTrap Heparin column using an Akta pure system. The column was washed with 10 column volumes (CV) of Buffer A and a gradient 0 to 100% Buffer B (pH7.5, 25 mM HEPES, 2 M NaCl) was applied over 10 CV, collecting 1 ml fractions.

The fractions were checked by western blot with anti-V5 (Dlp) and anti-Wg antibodies. GFP-Wg tends to elute at high salt, whereas GFP-Wg/Dlp elutes earlier in the gradient. The fractions containing GFP-Wg only were pooled and incubated with 1% CHAPS for 1 h at 4°C. They were then concentrated to 1 ml with Amicon^®^ Ultra Centrifugal Filter, 30 kDa MWCO (UFC8030) and loaded onto a Superdex^®^ 200 Increase 10/300 GL column (Cytiva 28-9909-44) equilibrated in pH7.5, 25 mM Hepes, 500 mM NaCl, 1% CHAPS. The run was performed at 0.5 ml/min in the above buffer, collecting 0.4 ml fractions. The fractions were run on a gel; the relevant fractions pooled, concentrated, snap-frozen, and stored at -80°C. Sample stability was determined using NanoDSF.

### GFP-Wg liposome and GFP-Wg aggregate generation

Following the protocol described in (Morrell et al., 2008), we diluted 1 μl of purified GFP-Wg (0.82 mg/ml in 25mM HEPES 150 mM NaCl 1% CHAPS) in 700 μl buffer without detergent (Buffer C, 25mM HEPES 150 mM NaCl) to make a working stock. 111 μl of this working stock were used to rehydrate 14 μmol of 1,2-Dimyristoyl-*sn*-Glycero-3-Phosphocholine (DMPC) (Avanti Polar Lipids 850345P) which had been freeze-dried from cyclohexane overnight. This solution was mixed with a magnetic stirrer at RT until all lipid had visibly dissolved. Buffer C was added to make a final volume of 250 μl and the solution was mixed and passed 31 times through a 100 nm filter using a Mini-Extruder (Avanti Polar Lipids) held at 35°C. The extruded solution was spun in an ultracentrifuge (Beckman Optima TLX, in a TLA100.3 rotor) at 43,000 rpm (100,000 g) for 30 min at 4°C. 32.5 μl samples of working stock, supernatant and pellet were saved, mixed with Bolt 4X sample buffer and 10X reducing agent (ThermoFisher B0007 and B0009) and boiled at 95°C for 5 min. The rest of the supernatant was discarded and the liposome pellet re-suspended in 250 μl Buffer C. A sample of liposomes was saved for the trypsin assay, and the rest was stored at 4°C for imaging the next day (see “Microscopy”).

To make GFP-Wg aggregates we used 1 μl of purified GFP-Wg (0.82 mg/ml) diluted as if to make liposomes (1:700 followed by 111:250) into Buffer C with or without 1% CHAPS. Samples were either immediately stored at 4°C or incubated for 1 h at RT before storage, and imaged the next day.

### Liposome trypsin assay

60 μl of resuspended liposomes were added to 420 μl DMEM (Gibco 41966-029) and 20 μl trypsin (Life technologies 25200-056) and incubated for 0 or 20 min at 37°C. The reaction was quenched with 1 ml of DMEM containing 10% fetal bovine serum (FBS). The samples were centrifuged at top speed for 30 min at 4°C, the supernatant was removed, and the pellet was resuspended in 60 μl DMEM and prepared for western blot as above.

### Western blot

Samples were run on 4-12% Bis-Tris Bolt gels (Invitrogen) with MES buffer (Invitrogen 13266499). Proteins on gel were transferred onto nitrocellulose membrane using Mini Blot Module transfer system (ThermoFisher). The membranes were washed with dH2O and blocked with 5% skimmed milk in TBST (TBS + 0.1% Tween-20) for 1 h at RT. Membranes were incubated with primary antibodies diluted in 5% milk TBST overnight at 4°C, and washed with TBST thrice before incubation with secondary antibodies for 2 h at RT. Membranes were washed with TBST thrice, incubated with 1:1 Clarity Max Western Blot ECL Substrate (Bio-Rad 1705062) for 5 min and imaged on a ChemiDoc MP imaging system (Bio-Rad). Primary antibodies used: mouse anti-Wingless (1:2000, DSHB 4D4), anti-V5 (1:4000 Abcam ab27671). Secondary antibodies used: anti-mouse HRP (1:2000, Jackson ImmunoResearch 715-035-151).

### Microscopy

Fixed wing discs were routinely imaged using a SP8 laser confocal microscope (Leica, DMI8-CS) controlled by Leica Application Suite X software (v. 3.5.5.19976). The objective used was HC PL APO CS2 63x/1.40 oil inmersion lens covered with Zeiss ImmersolT518 F oil (refractive index of 1.518). Bit depth resolution was set to 16, the scan direction was bidirectional, scan speed was 400 Hz, the pinhole size was set to 82.4 μm, and we used averaging of two lines with no frame accumulation/averaging. We used a white light laser set up for the following wavelengths, as required: 650 nm with HyD detector (658 – 800 nm), 558 nm with HyD detector (565 – 628 nm), 488 nm with HyD detector (498 – 540 nm); and a Diode 405 laser set up with a PMT detector (415 – 475 nm, gain 800). All HyD detectors were used in standard mode with gain set to 400. Laser powers, *xy* pixel size, *z*-stack size and *z*-step size were adjusted according to experimental needs.

Super-resolution imaging of fixed wing discs was performed using a SoRa CSU-W1 disk (Yokagawa), a 3.2× magnification lens (Olympus), and a photometrics BSI camera (95% QE, 6.5 μm pixels) mounted on an Olympus IX83 microscope. We used a 60x/1.5 NA UPLAPO OHR oil immersion lens covered with Olympus immersion oil type-F (refractive index of 1.518). Samples were excited with OBIS LS/LX 640-, 561-, 488-, 405 nm lasers (Coherent) (in that order), as required, with the corresponding emission bandpass filters: 685/40, 617/73, 525/50, 447/60. A variable number of *z*-sections was acquired to ensure the whole wing disc was captured, including peripodial cells for reference, but the thickness was kept constant at 0.3 μm per slice.

For live imaging of embryos, we used the same microscope set up with a few changes. We used a 60x/1.3 NA UPLSAPO S silicon immersion lens covered with Olympus silicon immersion oil (refractive index of 1.406) and because of the limitations of live imaging only thirteen *z-*sections with 0.5 μm in thickness each were acquired per image.

Liposomes and GFP-Wg aggregates were imaged on a similar SoRa CSU-W1 disk set up but with a Hamamatsu ORCA-Fusion camera (80% QE, 6.5 μm pixels) and a 60x/1.5 NA UPLAPO OHR oil immersion lens covered with Zeiss ImmersolT518 F oil. 30 ul of sample were added onto a 35 mm glass bottom Petri dish with a 20 mm micro-well (MatTek) pre-coated with poly-D-lysine (Cultrex, 3439-100-01) and a 1.5 round coverslip on top to protect samples (Epredia, 12 mm). Each sample was imaged in six areas chosen at random, and for each area we acquired a 20-slice *z*-stack with a *z*-step size of 0.24 μm.

For all SoRa microscopy, the controlling software was OLYMPUS cellSens Dimensions 4.1. This software was used to process the images using the automatic OSR enhancing filter post processing set to low followed by constrained iterative MLE deconvolution with five iterations and standard parameters. In all cases, laser power and exposure were adjusted according to the experiment.

To test if WgLVs could be detected using a different super-resolution technique that did not require deconvolution, SNAP-Wg wing discs were imaged also on a VT-iSIM microscope (VisiTech international) using an Olympus IX83 microscope, 150×/1.45 Apochromat objective (UAPON150XOTIRF) covered with Zeiss ImmersolT518 F oil and Prime BSI Express scientific CMOS camera (Teledyne Photometrics), controlled using Micro-Manager 2.0.0. We acquired a variable number of z sections to capture the whole wing disc with a 0.13 μm *z*-step size.

### Image analysis

#### Figure preparation

All images shown correspond to single *z-*slices unless otherwise stated. Apical *z*-slices were chosen as the apical-most slice were Wg signal was visibly in focus, basal *z-*slices were chosen as the most basal slice were Wg signal was still visible, lateral *z-*slices were chosen as the halfway point between the two of them for each wing disc. For each condition, representative images are shown, with at least five wing discs per condition analysed unless otherwise stated. Comparisons are only drawn between images acquired with the same laser power and exposure settings. Brightness and contrast levels were adjusted in Fiji (ImageJ2, release 2.17.0) but kept the same for all images being compared, except for maximum intensity projections in Fig7D (as the difference in background signal made this impossible). Example images of individual WgLVs were chosen at random from representative images and brightness/contrast settings were manually adjusted to keep local background and maximum intensity similar.

### WgLV quantification in super-resolution images

In images of GFP-Wg from live embryos, we quantified the size of WgLVs by manually finding them throughout the *z-*stacks and drawing a line to calculate an outer diameter. As this was extremely time-consuming, we wrote a semi-automated analysis pipeline to find small, bright ring-shaped objects in our wing disc images (from here, referred to us GOLLUM) using Fiji (ImageJ2, release 2.17.0), CellProfiler (version 4.2.5 (Stirling et al., 2021)) and R (The R Foundation for Statistical Computing, version 4.3.0) with RStudio (Posit Software PBC, 2025.05.0 Build 496). Examples of the scripts that make GOLLUM have been deposited on github, although some were tweaked to suit the different datasets (https://github.com/inesa-r/WgLV-Analysis-Gollum-).

First, the OSR processed and deconvolved image stacks were manually checked and corrected for chromatic aberrations due to sample thickness, then split into individual *z-*slices of the channel corresponding to Wg to be analysed with CellProfiler. On CellProfiler, WgLVs were identified by applying an Otsu filter with an experimentally adjusted size range requirement, and filtering the primary objects based on area, eccentricity and intensity distribution (heterogeneity, mean fractional intensity of central, 2^nd^ and overflow bins) according to experimentally adjusted parameters. This generated two masks per *z-*slice: one to identify WgLVs and a control of particles removed from the analysis. The performance of GOLLUM was compared to manual identification of WgLVs throughout two whole *z-*stacks of independent wing discs. “True positives” (TPs) were WgLVs found both by eye and by GOLLUM, “false negatives” (FNs) were WgLVs found by eye only, “GOLLUM positives” (GPs) had been missed by eye initially but were accepted as WgLVs, and “false positives” (FPs) were identified by GOLLUM but did not appear to fit the characteristics of WgLVs by eye. We then calculated the following scores as %:

Sensitivity by eye = (TPs+FNs)*100/(TPs+FNs+GPs)

Sensitivity by GOLLUM = (TPs+GPs)*100/(TPs+FNs+GPs)

Error rate by GOLLUM = (FPs)*100/(TPs+FPs+GPs)

Due to the highly variable difference in total Wg signal throughout the apical-basal axis, the GOLLUM pipeline was adjusted towards a medium performance, such that in the apical-most slices, GOLLUM sensitivity was almost as good as by eye, although the error rate was high; and in the lateral and basal slices, GOLLUM was more stringent, with a much lower error rate but missing many WgLVs (FigS1D). Given the extremely time-consuming nature of completely manual quantification, this was deemed an acceptable compromise when taking the disc as a whole, provided visual inspection of the images always accompanied the analysis, an additional quality-control via radial profiling to exclude sub-par wing discs (see below) and the caveat that identification of WgLVs on apical-most slices might not be as accurate.

All measurements of intensity were performed by referring to the equivalent position on the “raw” images (prior to super-resolution enhancement and deconvolution). The *z*-stacks were equally corrected for chromatic aberrations as their corresponding deconvolved *z*-stacks. If required, at this step we measured the mean intensity of the whole *z*-slice of a reference channel (e.g. anti-cadherin signal) to identify the most apical position. We also calculated a value of Wg intensity in Wg-producing cells over background applying an Otsu threshold to normalise the Wg intensity of WgLVs against, and that of any other test channels (e.g anti-Wls). We then applied the WgLV mask stack and the removed particles mask stack to the raw stack to find raw intensity values for each particle of interest in the deconvolved image, for each channel of interest. We also at this stage applied other masks if required, for example a mask for GMAP particles, a mask for Rab7 and/or Rab4, or a mask of the core of the DV boundary (Wg-producing cells) and measured the required parameters for the particles in those masks. The specific scripts to make those masks have been included in the github folder.

As mentioned, for extra quality control, we also used the two main mask stacks to calculate radial profiles with a maximum radius of 15 px of 40 particles of interest from each mask stack per wing disc, evenly distributed throughout the ROI list. We then mirrored the profiles and compared the average of all the radial profiles of WgLVs to that of all the radial profiles of removed particles. We expected one single central peak for the average of removed particles but two mirrored peaks further out for the WgLV average, if this was not the case, that wing disc was excluded from further analysis. We then calculated an average radial profile of all the wing discs that passed quality control. The same radial profile tool was used to analyse signal intensity distribution in WgLVs of other channels of interest (SNAP-surface649, YFP-Rab4 and YFP-Rab7). Note that because the radial profiles are normalised to their maxima for comparison, if there is no enrichment in WgLVs compared to the local background, the average is a flat line close to 1.0. Also because of averaging sometimes the two peaks corresponding to the WgLVs merged to form a wide peak, still noticeable different from the sharp peak of non-WgLV profiles.

Finally, we calculated for every disc that passed quality control the number of WgLVs and divided it by the number of slices in the stack to control for the number of images analysed. We also calculated the median size and median WgLV mean intensity of a channel of interest for all the WgLVs in that wing disc/compartment/mask as appropriate (median was chosen instead of average to avoid being skewed by outliers). The median of WgLV mean intensities was then divided by the median GMAP+ particles mean intensities to compare enrichment of Wg, GMAP or Wls in WgLVs versus Golgi (Fig1E) or normalised to the overall intensity over background calculated above to show general enrichment in WgLVs versus the cell as a whole.

### Extracellular Wg intensity quantification

All *z-*stacks were ordered apical to basal, selecting as the first (apical-most) *z-*slice the one corresponding to the transition from peripodial cells to the wing disc proper, and as last (basal-most) *z-*slice that were no nuclei were visible. A square of 25x25 pixels was placed either at the DV boundary or away from it (to measure the background). Mean intensity within the square was measured for all slices. The mean intensity of the background square was subtracted from the mean intensity of the DV square for each corresponding *z*-slice. The intensity over background was then plotted, using the number of slices and *z-*step size to calculate the distance from the apical-most slice.

### GFP-Wg aggregate quantification

To quantify the number and brightness of GFP-Wg aggregates, we manually thresholded the maximum intensity of every image (6 per condition) to select every possible foci from the background. We then calculated the average of the mean intensity of all the aggregates per image. The total GFP intensity in the image was the product of multiplying the total area covered by aggregates by the mean intensity of all the aggregates.

### Cell culture and luciferase assay

L-Wnt-3A cells (ATCC CRL-2647) were seeded in 12-well plates at a density of 5 × 10⁵ cells per well in DMEM supplemented with 10% FBS and incubated overnight at 37°C in a humidified incubator with 5% CO₂. The culture medium was then replaced with fresh DMEM containing FTY720 (Sigma, SML0700) at concentrations generated by two-fold serial dilution starting from 25 µM. Conditioned media were collected 7 days later and used for TOP-Flash luciferase assays.

STF293T cells (ATCC CRL-3249) were seeded into white-back 96-well plates at a density of 4 × 10⁴ cells per well in DMEM supplemented with 10% FBS. After 24 h, cells were transfected with the tk-Renilla luciferase plasmid (Promega; 30 ng per well) using TransIT transfection reagent (Merck), according to the manufacturer’s instructions. 24 hours post-transfection, the culture medium was replaced with L-Wnt-3A–conditioned medium prepared as described above. Firefly and Renilla luciferase activities were measured 24 h later using the Dual-Glo Luciferase Reporter Assay System (Promega) and an Ascent Luminoskan luminometer (Labsystems). Firefly luciferase activity was normalized to constitutive Renilla luciferase activity.

### Statistical analysis

No statistical methods were used to predetermine sample size. The experiments were not randomized, and investigators were not blinded to allocation during experiments and outcome assessment. Number of technical repeats, N and statistical test used are indicated in each figure legend, as necessary. Each dataset was tested for normal distribution using the D’Agostino–Pearson (omnibus K2) test. Exact *p-*values are reported, when possible, but to be as informative as possible without crowding the graphs, exact *p-*values are indicated only for all comparisons with *p* < 0.05 (Fig3D, 6C, 7E) or are summarised (Fig5D, FigS1H). Statistical analysis was performed in Prism 10 (GraphPad Software).

## Extended Data Tables

**Supplementary Table 1:** Fly lines used in this study.

| Fly line | Origin |
| --- | --- |
| *GFP-Wg* | (Port et al., 2014) |
| *GFP-Wg*<sup>S239A</sup> | (McGough et al., 2020) |
| *SNAP-Wg* | This study |
| *wls[ExGFP]* | (Yu et al., 2020) |
| *wls[ExHA]* | (generation described in (Yu et al., 2020), but not previously reported as such) |
| *YFP-Rab4* | (Dunst et al., 2015) |
| *YFP-Rab7* |  |
| *YFP-Rab5* |  |
| *YFP-RabX1* |  |
| *YFP-Rab8* |  |
| *YFP-Rab11* |  |
| *TagRFP-Vps35* | BDSC 66527 |
| *shibire*<sup>ts1</sup> | A gift from Cahir O’Kane, Cambridge University |
| *GFP-Chc* | This study |
| *nos-Cas9-mSA* | (Poernbacher et al., 2019) |
| *hs-Cre(w+)/Cyo* | (Siegal and Hartl, 2000) |
| *UAS-OptoCD8trap* | This study |
| *wg-Gal4* | (Alexandre et al., 2014) |
| *Ubi-Cry2olig-VHH-mCherry (III)* | This study |
| *naked-NeonGreen* | (Bolhaqueiro et al., under review) |
| *hh-Gal4* | (FlyBase, FBti0017278) |
| *UAS-schlank RNAi* | VDRC 33896 |
| *UAS-myr-RFP* | BDSC 7119 |
| *en-GAL4* | FlyBase, FBti0038265 |
| *UAS-RFP* | BDSC 30556 |
| *wg-Gal4DBD:2xnMagHigh1* | (di Pietro et al., 2021) |
| *wg[KO;attP]* | (Alexandre et al., 2014) |
| *wg-Gal80* | This study |
| *UAS-mCherry-nls; MKRS/TM6B* | BDSC 38425 |
| *hh[KO;attP]* | (Baena-Lopez et al., 2013) |
| *hh-Gal80* | This study |
| *schlank[KO;attP]* | (Sociale et al., 2018) |
| *FRT-schlank-HA-FRT* | This study |
| *vg-Flp* | (Bolhaqueiro et al., under review) |
| *UAS-GFP-Wg* | This study |
| *ubi-2xpMagHigh1:Gal4AD* | (di Pietro et al., 2021) |
| *en-Gal4, UAS-Flp* | (McGough et al., 2020) |
| *ubi-GFP, minute, FRT2A* | (Kakugawa et al., 2015) |
| *dlp[KO] (with and without FRT2A)* | (Bolhaqueiro et al., under review) |
| *dlp[KO; FRT-dlp-FRT]* | (Bolhaqueiro et al., under review) |
| *vg-Gal4* | (Stapornwongkul et al., 2020) |
| *UAS-Dlp-HA (WT)* | (Giráldez et al., 2002) |
| *UAS-Dlp*<sup>MN/CC</sup>-HA (Clamp) | (McGough et al., 2020) |

**Supplementary Table 2:**
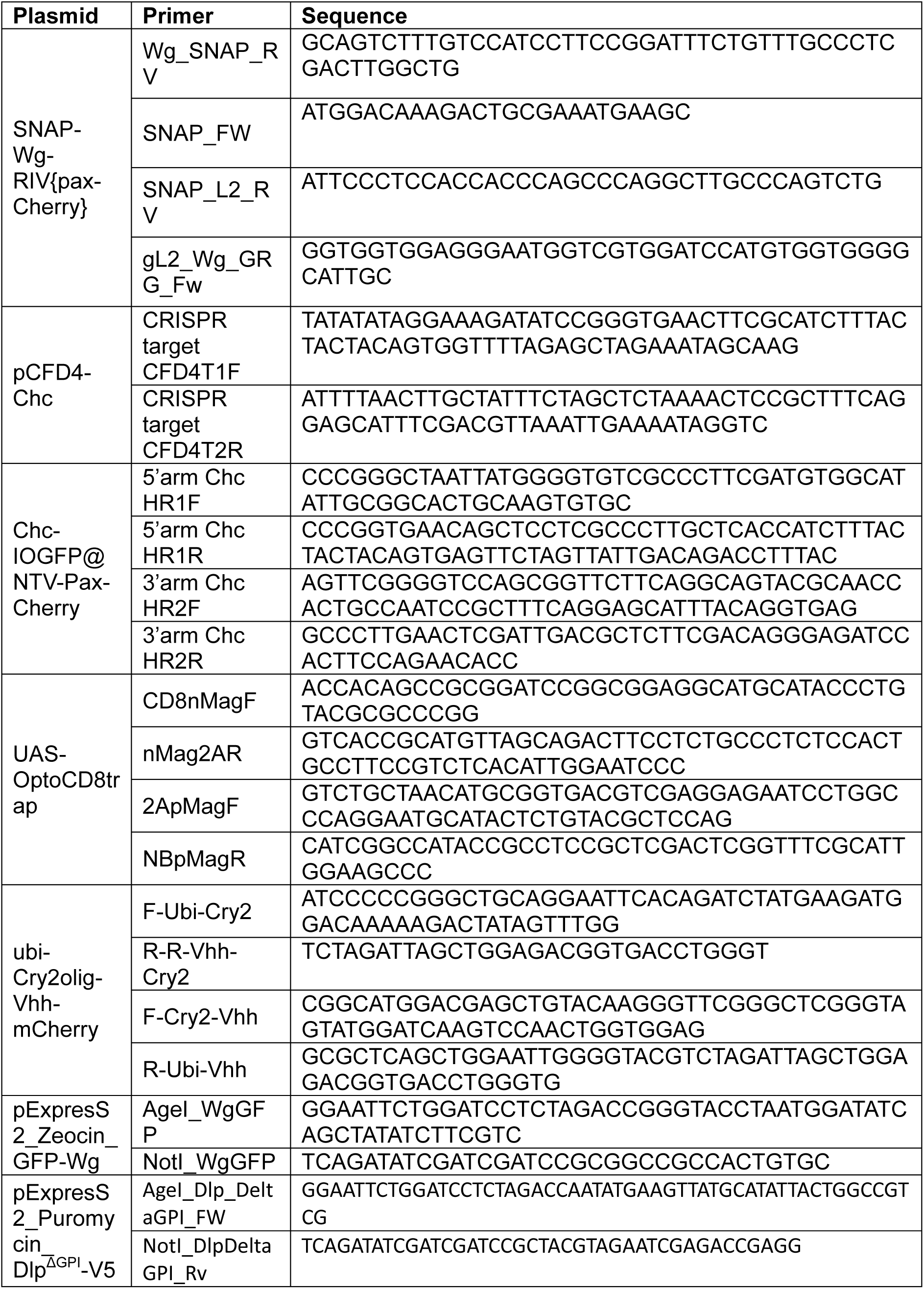
Sequences for fly line generation.

